# Integrated cellular and molecular responses to uranium chemotoxicity in the metal-tolerant microalga *Coelastrella* sp. PCV

**DOI:** 10.64898/2026.01.27.701972

**Authors:** Camille Beaulier, Fabienne Devime, Adrien Galeone, Grégory Si Larbi, Pierre-Henri Jouneau, Jonathan Przybyla-Toscano, Claude Alban, Stéphane Ravanel

**Affiliations:** Univ. Grenoble Alpes, INRAE, CNRS, CEA, IRIG, LPCV, 38000 Grenoble, France; Univ. Grenoble Alpes, CEA, IRIG, MEM, 38000 Grenoble, France

**Author notes:** Corresponding author Univ. Grenoble Alpes, INRAE, CNRS, CEA, IRIG, LPCV, 38000 Grenoble, France.

**Keywords:** Coelastrella, detoxification mechanisms, high-resolution imaging, ionomic, mineral homeostasis, tolerance, transcriptomic, uranium

## Abstract

Understanding the toxicity of hazardous metals in microalgae is critical for environmental risk assessment and sustainable phycoremediation. Metal-tolerant organisms provide powerful models for dissecting the mechanisms that mitigate metal toxicity. Here, we investigated the cellular and molecular responses to uranium (U) chemotoxicity in the metal-tolerant microalga *Coelastrella* sp. PCV. We used an integrated multi-omics and high-resolution imaging approach, combined with physiological analyses, to elucidate the mechanisms underlying U tolerance in *Coelastrella*. Using TEM-EDX, U was localized to the cell wall, polyphosphate bodies within acidocalcisomes, and vacuoles. Three-dimensional cell reconstruction and morphometric analysis using FIB-SEM showed that U-challenged cells displayed increased vacuolization, reflecting sequestration of uranyl ions and autophagy-mediated detoxification. Transcriptome responses were rapid and extensive, characterized by repression of cell division and photosynthesis, and pronounced imbalance in protein turnover and trafficking. Uranium also disrupted the homeostasis of essential elements, with marked rewiring of gene networks governing molybdenum, manganese, phosphate, iron and calcium homeostasis, notably affecting transporters and metal-binding proteins. *Coelastrella* sp. PCV efficiently sequestered U in acidocalcisomes and vacuoles, while rapidly excluding U from the cell. These coordinated detoxification responses are likely mediated by calcium, iron, ABC, and MATE transporters among the strongly deregulated genes under U stress.

## Introduction

Uranium (U) contamination of freshwater ecosystems is a persistent but often underestimated environmental threat (Markich, 2002). Although U is widely perceived through the lens of radiological risk, its primary mode of toxicity in biological systems is chemical (Gao *et al*., 2019). Green microalgae, the primary producers at the base of aquatic food webs, are directly impacted by hazardous metal exposure, threatening ecosystem functioning, carbon fixation, and water quality. Understanding the chemical toxicity of U in these organisms is therefore essential for environmental risk assessment and the development of sustainable phycoremediation strategies.

The concentration of U in surface freshwaters is governed by both geochemical context and anthropogenic activities. In oxic aquatic environments, U occurs predominantly as hexavalent U(VI), in the form of free or complexed uranyl ions (UO_2_^2+^) depending on water physicochemical properties (Markich, 2002). Uranium bioavailability is primarily controlled by chemical speciation, which depends on pH, carbonate and phosphate concentrations, water hardness, and dissolved organic carbon (e.g. (Franklin *et al*., 2000; Charles *et al*., 2002; Hogan *et al*., 2005; Fortin *et al*., 2007)). At the molecular level, U toxicity is driven by its strong coordination chemistry, resulting in U(VI) binding to phosphate, carboxyl, and carbonate groups of biomolecules. Green microalgae possess a remarkable capacity to bind U at the cell surface through a rapid and passive biosorption process, removing a large fraction of dissolved U within minutes and concentrating the element at the cell-environment interface where desorption, biomineralization or uptake may subsequently occur. Spectroscopic studies have shown that carboxyl, amino and phosphate groups are involved in U complexation to algal cell walls (Gunther *et al*., 2008; Vogel *et al*., 2010). Evidence for intracellular U accumulation remains limited, but electron-dense inclusions containing U have been detected in the cytoplasm of *Chlamydomonas* sp. (Garcia-Balboa *et al*., 2013). Consistent with U internalization, cell division and photosynthesis emerged as primary targets of U chemical toxicity across diverse algal species, with exposure rapidly impairing cell proliferation (e.g. (Franklin *et al*., 2000; Charles *et al*., 2002; Hogan *et al*., 2005)) and photosynthetic performance (Herlory *et al*., 2013).

Most studies of U toxicity have historically focused on laboratory strains such as *Chlamydomonas reinhardtii* and *Chlorella vulgaris*, until the identification of U-tolerant green algae isolated from contaminated environments (Baselga-Cervera *et al*., 2018; Beaulier *et al*., 2024). These organisms revealed that tolerance may involve active and dynamic coping strategies. Recent work on *Coelastrella* sp. PCV exemplifies this behavior, demonstrating exceptional tolerance to U concentrations that are lethal to standard freshwater models while maintaining photosynthetic activity and growth (Beaulier *et al*., 2024). Notably, this microalga rapidly sequesters U from the medium, followed by delayed release, effectively functioning as a U excluder. Such biphasic behavior suggests regulated desorption and/or efflux mechanisms that mitigate chemical toxicity.

At the molecular level, U responses in the green lineage are far better understood in land plants than in microalgae. Uranium uptake has been linked to calcium(Ca)-permeable channels in *Arabidopsis thaliana* root cells (Sarthou *et al*., 2022) and to endocytic processes in *Nicotiana tabacum* cells (John *et al*., 2022). Once internalized, U interacts with phosphate-containing metabolites (Berthet *et al*., 2018) and proteins (Sarthou *et al*., 2020). Recent works have identified U-binding proteins potentially involved in metal sequestration or contributing to toxicity, for example through interference with RNA metabolism (Vallet *et al*., 2023; Revel *et al*., 2025). Transcriptomic and proteomic analyses in plants further revealed coordinated stress responses, together with deregulation of cell wall biogenesis, hormone signaling pathways, and iron and phosphate homeostasis under U exposure (Doustaly *et al*., 2014; Lai *et al*., 2020; Chen *et al*., 2025; Przybyla-Toscano *et al*., 2025). By contrast, the molecular mechanisms underlying U stress responses, tolerance, and dynamic accumulation remain largely unknown in green microalgae. Given its unique response to U, *Coelastrella* sp. PCV provides a compelling model to address this knowledge gap. In the present study, we integrated high resolution cell and metal imaging with genomics, transcriptomics and ionomics to dissect responses to U chemotoxicity and to identify candidate mechanisms driving U tolerance and accumulation in this metal-tolerant microalga.

## Materials and methods

### Culture conditions

*Coelastrella* sp. PCV was cultivated as described in (Beaulier *et al*., 2024). Cells were maintained in TAP medium at 21°C in continuous white light (40 µmol photons s^-1^.m^-2^). Uranium stress was applied in a modified low-phosphate TAP medium (LoP, pH 7.0) containing a reduced phosphate concentration (50 µM instead of 1 mM) to limit the formation of U-phosphate precipitates. Exponentially growing cells in TAP were transferred at a final density of 1 million cells/mL to LoP supplemented or not with 200 µM uranyl nitrate (U200) and growth was pursued for up to 24 or 48 h. Growth rates were assessed by cell counting using a Luna automated cell counter (Logos biosystems). Photosynthetic parameters were measured using a Maxi-Imaging PAM fluorometer (Heinz Walz), as described in (Beaulier *et al*., 2024). Biological replicates are defined as samples obtained from separate culture flasks, grown and processed independently.

### Sample preparation, sequencing, and *de novo* nuclear genome assembly

The extraction of high-molecular-weight and high-purity genomic DNA (gDNA) from *Coelastrella* sp. PCV was performed using a procedure adapted from (Stark *et al*., 2020). Cells growing exponentially in TAP medium were harvested by centrifugation, suspended in extraction buffer (50 mM Tris, pH 8.0, 20 mM EDTA, 350 mM NaCl, 2% w/v SDS), flash-frozen and homogenized by grinding in liquid nitrogen. After thawing, one volume of CTAB buffer (50 mM Tris, pH 8.0, 20 mM EDTA, 1.4 M NaCl, 2% w/v CTAB) was added to homogenized cells and incubation was pursued for 1 h at 60°C. Nucleic acids were treated with RNase A, extracted with phenol/chloroform/isoamyl alcohol and chloroform, precipitated with ethanol, and washed with 70% v/v ethanol. Genomic DNA was dissolved in TE and the quality of the preparation was assessed using an Agilent 2200 TapeStation, which demonstrated a high-molecular-weight and high-quality DNA (DIN value of 9.3). Genomic DNA was used for both long-read PacBio and short-read Illumina sequencing technologies. First, a PacBio Sequel II SMRTbell library was prepared for high-fidelity sequencing on a PacBio Sequel II platform (Macrogen Europe). Second, an Illumina TruSeq DNA PCR-free library was prepared for sequencing on an Illumina NovaSeq 6000 platform (Macrogen Europe). The PacBio Sequel II was configured to produce up to 15 Gb of high-fidelity reads with insert sizes up to 15 kb, while the NovaSeq 6000 was set for 150 bp paired-end sequencing. Raw sequencing data statistics are summarized in Table S1.

Total RNA was purified from Coelastrella cells grown exponentially in TAP medium using the RNeasy Plant Mini Kit (Qiagen). The RNA integrity was assessed and validated using an Agilent 2200 TapeStation (RIN value of 9). For RNA sequencing, an Illumina TruSeq Stranded mRNA library was prepared and sequenced on a NovaSeq 6000 system set to generate 30 M of 150 bp paired-end reads. Raw Illumina reads were analyzed with FastQC (www.bioinformatics.babraham.ac.uk/projects/ fastqc/) and trimmed using BBDuk v38.75 (Bushnell *et al*., 2017), with minimum quality score set to 25 and minimum read length set to 35 bp, resulting in 97,875,342 trimmed read pairs for DNA sequencing and 71,857,762 trimmed read pairs for RNA sequencing. Trimmed Illumina DNA reads were analyzed using Genomescope (http://genomescope.org/genomescope2.0/) to estimate genome size based on 21 K-mer distribution. The estimated genome size was 113.44 million bp with a heterozygosity estimation of 0.044%, indicating a highly homozygous or haploid sample.

The PacBio sequencing produced 5,785,864 reads with an average length of 65.69 kbp and a N50 of 145.211 kbp. *De novo* genome assembly from PacBio data was conducted using CANU (Koren *et al*., 2017). Assembly polishing was performed with Pilon (Walker *et al*., 2014) and Racon (https://github.com/lbcb-sci/racon) by incorporating the Illumina short-read data. The assembly was evaluated using QUAST (https://sourceforge.net/projects/quast/), and the key statistics are summarized in Table S2. The completeness of the genome assembly was evaluated using BUSCO (Seppey *et al*., 2019) against the databases of Eukaryota and Chlorophyta conserved genes.

### Nuclear genome annotation

Nuclear genome annotation was conducted using the MAKER pipeline (Campbell *et al*., 2014), including a GeneMark (Lukashin & Borodovsky, 1998) prediction model supported by RNA-seq mapped reads, an Augustus (https://github.com/Gaius-Augustus/Augustus) prediction model generated from the best genes predicted by GeneMark, and homology-based evidence from Chlorophyta UniRef protein sequences. RNA-seq reads were aligned using STAR v2.7.3a (https://github.com/alexdobin/STAR) in double pass mode, achieving a unique mapping rate of 92.3%. The genome-guided transcriptome assembly was performed using Trinity (https://github.com/trinityrnaseq). Transcript redundancy was reduced using cd-hit-est (Li & Godzik, 2006), and assembly artifacts were filtered out using Kallisto v0.46.0 (Bray *et al*., 2016), which applies pseudoalignment to estimate expression levels. To refine gene models, an additional annotation pass was performed with StringTie (Shumate *et al*., 2022). Functional annotation was assigned using PANNZER2 (Törönen *et al*., 2018) and Trinotate (https://github.com/Trinotate) using a database of UniProtKB Viridiplantae proteins. Ribosomal RNA genes were annotated using Barrnap (https://github.com/tseemann/barrnap).

### Chloroplast and mitochondrial genomes assembly and annotation

To perform a specific assembly and annotation of chloroplast and mitochondrial genomes, the Illumina and PacBio reads were mapped against the *Coelastrella vacuolata* sequences (Shetty *et al*., 2021). Using the minimap2 algorithm (Li, 2018), about 545,000 Illumina and 108,000 PacBio reads could be mapped to the Cp genome, while about 16,000 Illumina and 16,000 PacBio reads were mapped to the Mt genome. For the cpDNA genome, the best assembly was obtained using Unicycler (https://github.com/rrwick/Unicycler) with the bold option activate, followed by a refinement with CAP3 (Huang & Madan, 1999), and finally a polishing step composed by 5 iterations of Racon (PacBio long-read polisher) and 10 iterations of Pilon (Illumina short-read polisher). The resulting assembly was composed by one contig of 213,460 bp. The annotation was performed with the GeSeq online service (Tillich *et al*., 2017) selecting *Coelastrella saipanensis*, *Pectinodesmus pectinatus* and *Tetradesmus obliquus* as references. For the mtDNA genome, the best assembly was obtained using wtdbg2 (Ruan & Li, 2020) followed by a polishing step composed by 10 iterations of Racon and 10 iterations of Pilon. The resulting assembly was composed of one contig of 69,229 bp. Due to the low coverage of the Illumina data (less than 80% of the genome being covered by at least one read), the assembly relied primarily on the PacBio data. The annotation was performed with the GeSeq online service selecting *Pectinodesmus pectinatus* as reference (Zhao *et al*., 2022). The structure and annotation of the cpDNA and mtDNA were produced with OGDRAW (Greiner *et al*., 2019).

### RNA preparation and sequencing

Erlenmeyer flasks containing 20 mL of LoP medium, with or without 200 µM uranyl nitrate (U200), were inoculated at 1 million cells/mL. Cells were collected by centrifugation (6,000 g, 3 min) after 0, 0.5, 1, 4, and 24 h of culture, in quadruplicate, and snap-frozen in liquid nitrogen. Frozen cell pellets were resuspended in 350 µL RNAprotect Tissue Reagent (Qiagen) and incubated overnight at 4°C. RNAprotect was removed by centrifugation (5,000 g, 2 min) and total RNA was purified using the RNeasy Plant Mini Kit (Qiagen), including DNase I treatment. RNA quality determined using an Agilent 2200 TapeStation provided RIN values above 7 for all RNA samples. For RNA-seq, TruSeq Stranded Total RNA libraries from Ribo-Zero Plus–depleted RNA (Illumina) were prepared and sequenced on a NovaSeq 6000, generating 30 million 150-bp paired-end reads.

### Differential expression analysis

RNA-seq reads were mapped to the reference *Coelastrella* sp. PCV genome using STAR (version 2.7.9a, https://github.com/alexdobin/STAR). FeatureCounts (version 2.0.0) was used to calculate gene expression values as raw fragment counts. The ARSyNseq function in the NOISeq R package was then used to filter raw fragment counts after TMM normalization (Nueda *et al*., 2012). Last, the HTSFilter package (https://github.com/andreamrau/HTSFilter) was used to remove genes with low and constant expression levels. Differentially expressed genes (DEGs) were defined as having a minimal expression of 1 fragment per kilobase million (FPKM) in at least one sample in the experiment, a |logfold-change| ≥ 1 in at least one treated (U200) vs control (LoP) comparison, and a Benjamini-Hochberg adjusted p-value <0.01.

### qPCR analysis

qPCR analysis was performed using RNAs purified for the RNA-seq experiment. Total RNA (1 µg) was reverse transcribed using the QuantiTect reverse transcription kit (Qiagen). qPCR reactions were performed using the PowerSYBR Green PCR mastermix (Applied Biosystems) on a CFX Connect Real-Time PCR System (BioRad). Gene expression was quantified using the ΔΔCt method, with G5680 (actin) used as the reference gene and time point 0 in LoP as the control condition (CFX Maestro v1.1, BioRad). Primers sequences of target genes are listed in Table S3.

### Inductively coupled plasma mass spectrometry

Cells were separated from the medium by centrifugation (4 min at 10,000 g), washed three times with 10 mM sodium carbonate, and dried at 80°C. Dried samples were digested in 65% (w/v) HNOfor 2 hours at 90°C. Cell and medium samples were diluted to the appropriate concentration and analyzed using an iCAP RQ quadrupole mass spectrometer (Thermo Fisher Scientific GmbH, Germany). Elements were analyzed using the standard (for 24Mg, 25Mg, 31P, 39K, 55Mn, 95Mo, 98Mo, 238U) and the collision mode with helium as a cell gas (for 31P, 39K, 44Ca, 55Mn, 56Fe, 57Fe, 63Cu, 65Cu, 64Zn, 66Zn). Quantification was done using standard curves and internal standards (45Sc, 103Rh, 172Yb).

### Scanning electron microscopy

Algal cells in TAP medium were deposited as a droplet onto Whatman paper, then supplemented with a droplet of 2% (v/v) ionic liquid (Hitachi HILEM IL1000; ethyl(2-hydroxyethyl)dimethylammonium methanesulfonate). Freshly prepared samples were imaged by scanning electron microscopy using a Quanta FEG-250 microscope (FEI, USA) operating at 2 kV under high vacuum conditions.

### Scanning transmission electron microscopy and energy dispersive X-ray analysis

*Coelastrella* cells were harvested by centrifugation, washed once in LoP medium, cryofixed using high-pressure freezing (210 MPa and -196 °C; EM HPM100, Leica, Germany) followed by freeze substitution (EM ASF2, Leica, Germany), as described in (Gallet *et al*., 2024). Uranyl acetate was omitted from the procedure to avoid interference with U detection in stressed cells. Ultrathin sections (70–80 nm) were prepared from resin-embedded samples using a diamond knife on a PowerTome ultramicrotome (Boeckeler Instruments, USA) and collected on 200-mesh copper grids. Imaging was performed by scanning transmission electron microscopy on a Tecnai microscope (FEI, USA) set up at 200 kV combined with energy dispersive X-ray spectrometer (Bruker Corporation, USA).

### Focused ion beam - scanning electron microscopy imaging

Focused ion beam (FIB) tomography was performed on a Zeiss CrossBeam 550 microscope (Zeiss), equipped with Fibics Atlas 3D software for automated tomography, as described in (Uwizeye *et al*., 2021). Resin blocks containing embedded cells were metallized with a 4 nm platinum layer to avoid charging during observations. Inside the FIB-SEM, a second platinum layer (1–2 µm) was locally deposited on the region of interest to mitigate curtaining artefacts. Serial ablation was then performed using a Ga^+^ ion beam (usually 700 nA at 30 kV). After each milling step, the freshly exposed surface was imaged by scanning electron microscopy at 1.5 kV with a current of c.a. 1 nA using the in-column EsB backscatter detector. The simultaneous milling and imaging mode was used to improve stability, with automated hourly correction of focus and astigmatism. A slice thickness of 8 nm was removed at each cycle, and SEM images were recorded with a pixel size of 8 nm, resulting in an isotropic voxel size of 8 × 8 × 8 nm^3^. Aligned FIB-SEM stacks were cropped using the Fiji software (https://imagej.net/Fiji) to isolate the region of interest. Three-dimensional segmentation, reconstruction, and quantitative analyses (surface area and volume) were performed using Dragonfly software (www.theobjects.com/dragonfly) (Uwizeye *et al*., 2021; Ezzedine *et al*., 2023). The Blender software (https://www.blender.org/) was used to capture 2D images of the cell ultrastructure from 3D reconstructions.

### Staining of vacuoles using Cell Tracker Blue

Cells were washed in LoP and incubated for 45 min in darkness with 6 µM Cell Tracker Blue CMAC (7-amino-4-chloromethylcoumarin; ThermoFisher Scientific). Stained cells were pelleted by centrifugation, resuspended in LoP, and imaged with an Axioplan 2 microscope (Zeiss) using the DAPI fluorescence filter (excitation 365, emission 445/50).

### Immunoblot analysis

Total proteins were extracted in 20 mM Tris pH 7.5, 1 mM EDTA, 2% (w/v) SDS, protease inhibitors and clarified by centrifugation (20,000 g, 15 min, 4°C). Proteins were separated by SDS–PAGE, transferred to nitrocellulose, and probed with antibodies against chloroplastic Fe-SOD (AS06 125, Agrisera), catalase (AS15 2991, Agrisera), ATPB (AS03 030, Agrisera) and PsaD (AS09 461, Agrisera). Fluorescence signal of Alexa Fluor Plus 488-conjugated secondary antibodies (Invitrogen) was detected on a Typhoon RGB imager (Cytiva) using a 488-nm laser and a Cy2 filter.

### Accession numbers

Genomic and RNA sequencing data have been deposited to the NCBI Sequence Read Archive under BioProject PRJNA1219995.

## Results and discussion

### 1 *Coelastrella* sp. PCV *de novo* genome assembly and annotation

High-quality genomic and transcriptomic datasets were obtained using the integration of long-read PacBio and short-read Illumina sequencing of both genomic DNA and RNA from *Coelastrella* sp. PCV cells growing exponentially in TAP medium. The final *de novo* nuclear genome assembly comprised 407 contigs with a total size of 119.8 Mb and a GC content of 51.71% (Table S2). The assembly had a N50 of 5.97 Mb and a L50 of 8, indicating a well-assembled and contiguous genome. 93.36% of the Illumina DNA reads mapped back to the assembly, supporting its accuracy. BUSCO analysis revealed 92.2% completeness with Eukaryota markers and 93.2% with Chlorophyta markers (Fig. S1), confirming the high-quality, near-complete assembly of the *Coelastrella* sp. PCV nuclear genome. Mapping of RNA-seq reads showed 92.3% unique alignments, ensuring high reliability for downstream expression analysis.

Annotation of the *Coelastrella* sp. PCV nuclear genome identified 15,856 genes and 16,802 protein-coding transcripts. This high-quality assembly is substantially larger and encodes an expanded gene repertoire compared to previously available *Coelastrella* draft genomes, which range from 75 to 83 Mb with 3,803 to 11,162 predicted genes (Karpagam *et al*., 2018; Shetty *et al*., 2021; Baldanta *et al*., 2025). Both the genome size and gene content of *Coelastrella* sp. PCV are comparable to those reported for microalgae of the Scenedesmaceae family, spanning 101 to 151 Mb and encoding 12,000 to 17,500 genes (Knoshaug *et al*., 2020; Calhoun *et al*., 2021; Biondi *et al*., 2024). The chloroplast DNA of *Coelastrella* sp. PCV was assembled into a unique contig resulting in a circular genome of 213,460 bp (Fig. S2). The annotation identified 31 tRNA, 24 rRNA, and 87 protein-coding genes mainly associated with the photosynthetic machinery, transcription, and translation. The *Coelastrella* sp. PCV plastome is comparable to those of *Coelastrella vacuolata* MACC-549 and other members of the Sphaeropleales order in terms of size and gene content (Shetty *et al*., 2021; Zhao *et al*., 2022). The assembly of the mitochondrial genome resulted in a unique contig of 69,229 bp (Fig. S2). A total of 35 tRNA, 3 rRNA, and 13 protein-coding genes were annotated, all linked to the mitochondrial respiratory function. The size and gene repertoire of the mitochondrial genome are more variable among *Coelastrella* sp. PCV, Sphaeropleales, and other green microalgae than those of the chloroplast genome (Shetty *et al*., 2021; Zhao *et al*., 2022). The diversity in gene organization and architecture within mitochondrial genomes has already been described as a puzzling aspect of the molecular evolution of green algae (Leliaert *et al*., 2012).

### 2 Cellular architecture of *Coelastrella* and consequences of U stress

*Coelastrella* sp. PCV cells growing exponentially in mixotrophic conditions (TAP medium) were imaged by scanning electron microscopy (SEM). Cell shape and morphology were strongly influenced by the sample preparation procedure (e.g. fixation with glutaraldehyde, dehydration), resulting in varying degrees of surface wrinkling and irregularity that do not reflect the native state of the alga observed by optical microscopy (Beaulier *et al*., 2024). To minimize artifacts due to chemical treatments, freshly harvested cells were treated with 2% (v/v) ionic liquid and observed by SEM. Images showed round-shaped cells with irregular surfaces characterized by numerous faint wrinkles and well-defined meridional ribs (Fig. 1A). Similar characteristics have been reported in various *Coelastrella* species (Wang *et al*., 2019; Kawasaki *et al*., 2020). Methodological artifacts affecting cell morphology require alternative approaches for reliable structural analysis. To this aim, we further analyzed the three-dimensional subcellular architecture of *Coelastrella* sp. PCV using focused-ion-beam scanning electron microscopy (FIB-SEM). Cells grown in TAP were cryo-fixed by high-pressure freezing, followed by slow freeze substitution and resin embedding, to maximize preservation of native structures (Uwizeye *et al*., 2021; Ezzedine *et al*., 2023). FIB-SEM datasets were processed to generate 3D reconstructed cell models suitable for quantitative morphometry of (sub)cellular structures. Ultrastructure analysis of multiple cells from the stack revealed no significant variation in the general organization of individual *Coelastrella* cells. The cell wall surface was relatively smooth (Fig. 1B), confirming that the observation of ribs in SEM images is strongly influenced by the sample preparation method. The cell wall thickness measured at the cell meridian was 90 ± 12 nm (n=5 individual cells), consistent with previously reported values for a Nordic *Coelastrella* species (Spain & Funk, 2022).

**Fig. 1:**
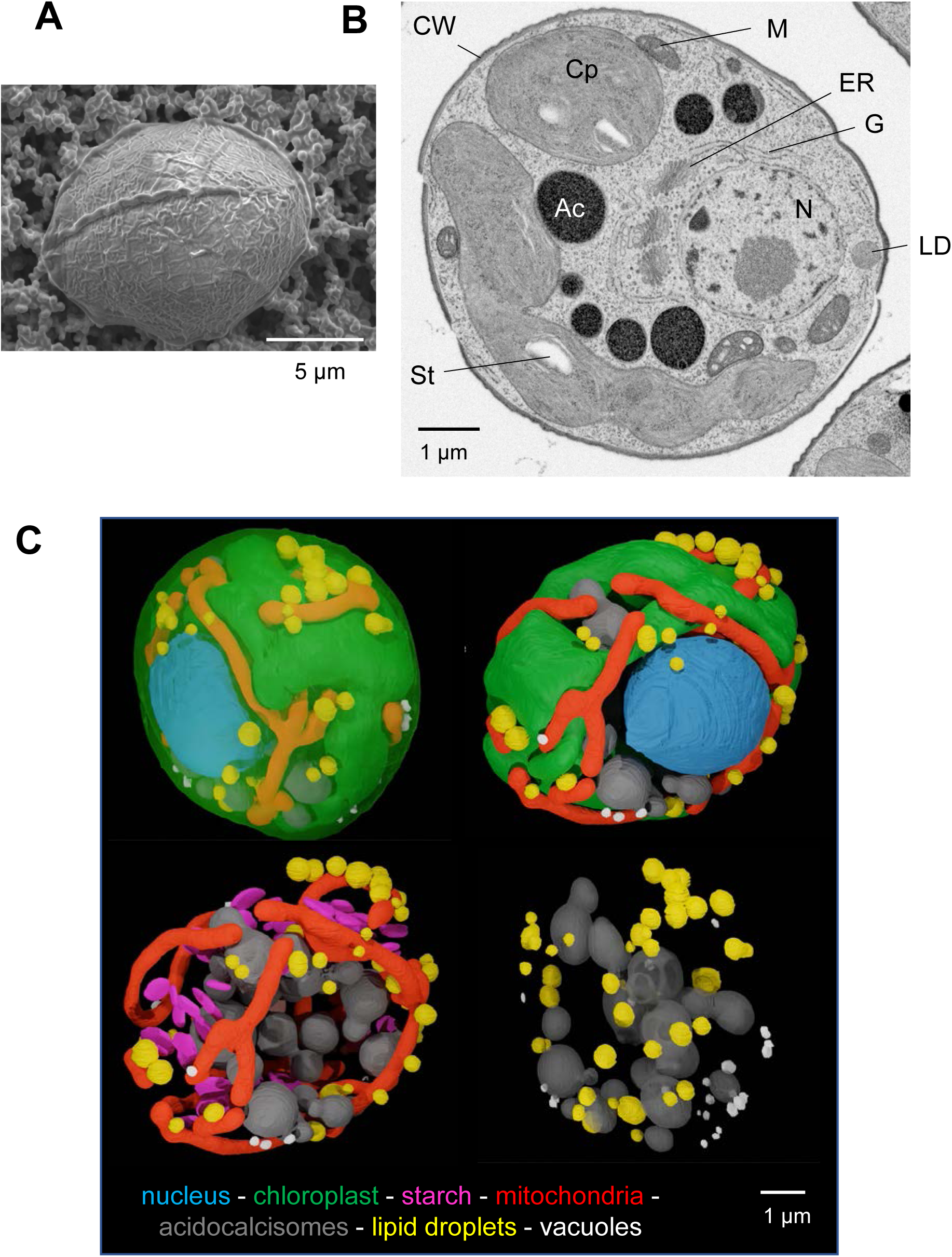
Cellular and subcellular architecture of *Coelastrella* sp. PCV. Cells were grown in TAP medium at 21°C and 40 µmol of photons.m^-2^.s^-1^ in continuous light. **A** - SEM micrograph of a native *Coelastrella* cell deposited on Whatman paper and treated with 2% (v/v) ionic liquid. **B** - FIB-SEM micrograph of a cryo-fixed and cryo-substituted *Coelastrella* cell. Ac, acidocalcisome; Cp, chloroplast; CW, cell wall; ER, endoplasmic reticulum; G, Golgi; LD, lipid droplet; Mt, mitochondrion; N, nucleus; St, starch granule. **C** - 3D reconstructed *Coelastrella* cell. Segmentation of FIB-SEM images highlighted the main subcellular compartments: chloroplast in green, mitochondria in red, nucleus in blue, acidocalcisomes in grey, lipid droplets in yellow, and vacuoles in white. Images are cropped from Movie S1.

Two *Coelastrella* cells grown in TAP were segmented and reconstructed in 3D. A representative cell contains a unique chloroplast occupying 36% of the cell volume and ten tubular elongated mitochondria (3-4% of the cell volume) that are intricately enclosed within chloroplast invaginations (Fig. 1C; Table S4; Movie S1). Other subcellular structures that could be quantified using the FIB-SEM tomography workflow include the nucleus (6% of the cell volume), large electron-dense and almost spherical structures (6%), lipid droplets (0.3-1%), and small electron-lucent vacuoles (0.1%) (Table S4). Electron-dense spherical structures were defined as acidocalcisomes containing polyphosphate (polyP) granules by TEM coupled to energy-dispersive X-ray spectroscopy (EDX). Indeed, the EDX spectrum of these granules indicated the presence of phosphorus and Ca (Fig. S3), which is the elemental signature of these lysosome-related acidic organelles involved in ion storage, osmoregulation, and pH homeostasis (Hong-Hermesdorf *et al*., 2014; Docampo, 2016; Tsednee *et al*., 2019; Schmollinger *et al*., 2021). The morphometric parameters determined for *Coelastrella* sp. PCV are comparable to those observed for various unicellular algae, in which the nuclei, plastid(s) and mitochondria fill 40-55% of the total cell volume (Uwizeye *et al*., 2021). In *Chlamydomonas* species, acidocalcisomes have been characterized as highly variable in number and size, occupying about 1% of total cell volume (Siderius *et al*., 1996; Ruiz *et al*., 2001). In *Coelastrella* sp. PCV grown in nutrient-replete conditions (TAP medium), the number of acidocalcisomes was also varying importantly (from 9 to 28, with an average of 18) and the cell volume occupancy (4.2 ± 0.7%, n=7 cells) was substantially higher than in *Chlamydomonas* (Fig. S4).

To analyze the consequences of U stress on the morphology and cellular organization of *Coelastrella* sp. PCV, cells were challenged for 24 h with 200 µM uranyl nitrate (U200) in a modified TAP medium with low phosphate content (50 µM instead of 1 mM, LoP medium). Under these conditions, U bioavailability is moderately affected by the precipitation of U-phosphate complexes and toxic effects are observable, i.e. reduced growth and photosynthesis (Beaulier *et al*., 2024). Cells growing in TAP were transferred to LoP or U200 for 24h, rinsed once with LoP, cryo-fixed and cryo-substituted, and analyzed by FIB-SEM. The ultrastructure of *Coelastrella* cells appeared consistent across multiple cells analyzed from both the LoP and U200 stacks, demonstrating structural consistency in both conditions. Two representative cells grown in either LoP or U200 were segmented, reconstructed in 3D, and used for morphometric analysis (Table S4; Fig. 2). *Coelastrella* cells from the LoP and U200 conditions shared many characteristics, of which the cell volume occupied by the chloroplast (30-38%), mitochondria (1.7-2.6%), the nucleus (3.8-5.5%), and acidocalcisomes (9.9-14.7%) (Table S4).

**Fig. 2:**
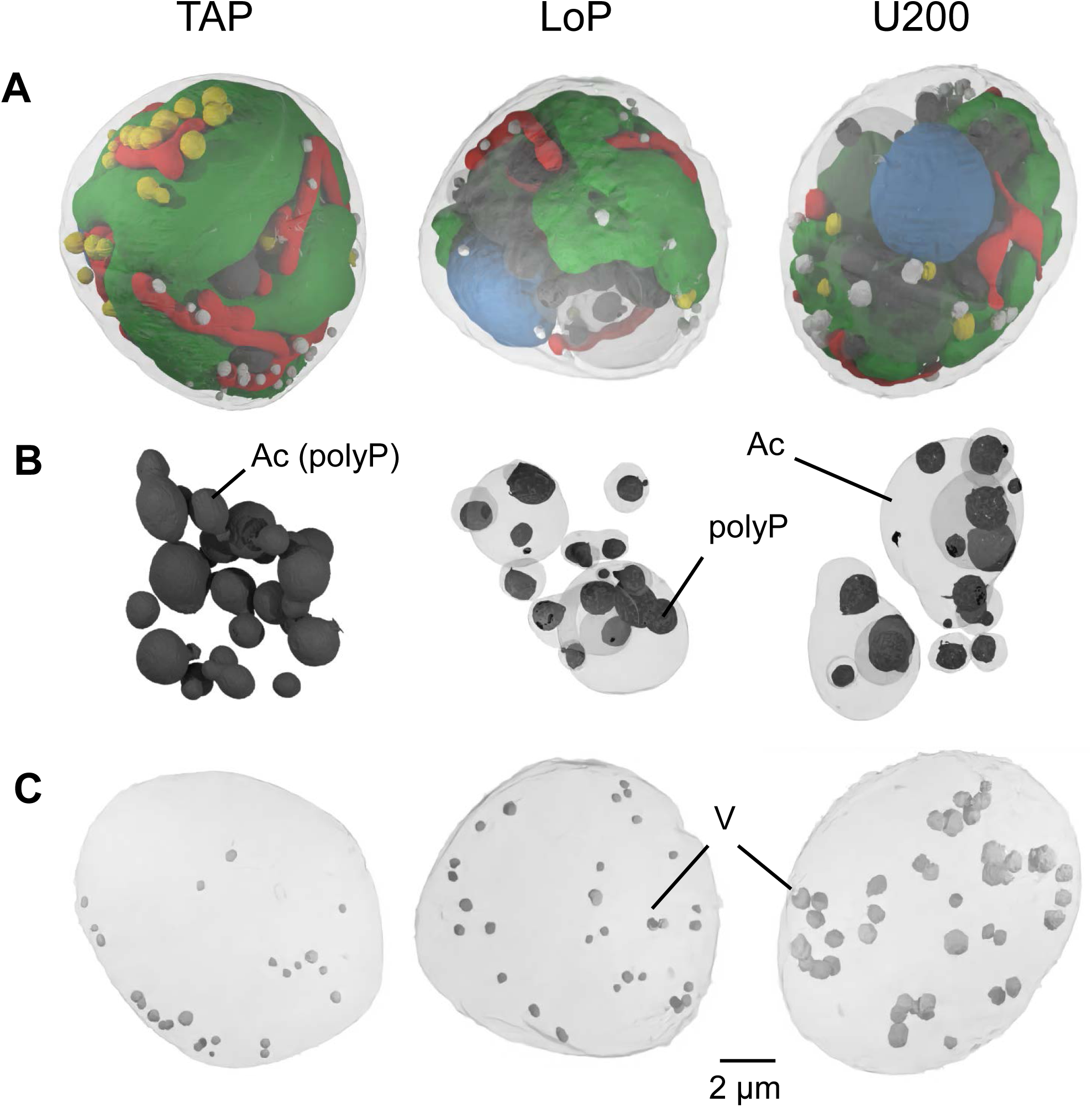
Morphological changes of acidocalcisomes, polyP granules and vacuoles in *Coelastrella* challenged with U. Cells were grown for 24 h in either TAP, LoP, or U200, vitrified and analyzed using FIB-SEM. **A** – 3D reconstructed cells showing chloroplasts in green, mitochondria in red, nucleus in blue, and lipid droplets in yellow. **B** –Structural changes in acidocalcisomes (Ac), including their polyP granules (polyP). **C** – Structural changes in vacuoles (V).

Significant changes in the ultrastructure and cell volume occupancy of acidocalcisomes were observed in LoP and U200 cells as compared with TAP (Fig. 2, S5). First, although the number of acidocalcisomes remained comparable, their volume fraction was increased by 2-fold in LoP and U200 conditions compared with TAP (Table S4). This difference was confirmed through quantitative measurements performed on several cells from the respective stacks (cell volume occupancy of 4.2 ± 0.7%, 10.2 ± 3.3%, and 9.7 ± 2.9% in TAP, LoP, and U200, respectively; n=6-7 cells) (Fig. S4). Also, FIB-SEM and TEM-EDX analysis revealed acidocalcisomes with varying electron-dense core patterns, from fully filled electron-dense matrix in TAP to partially filled structures in LoP and U200 (Fig. 2, S5). The electron-dense matrix made of polyP bodies (Docampo, 2016; Augusto *et al*., 2025) occupied the entire acidocalcisome volume in TAP, compared with 6-12% in LoP and U200 (Table S4, Fig. S4). The effects of phosphate limitation (LoP) and U stress (U200 in LoP) on acidocalcisomes were undistinguishable, suggesting that phosphate limitation is the main driver of these ultrastructural changes. Indeed, studies on *C. reinhardtii* grown under varying phosphate regimes (depletion and repletion) showed that polyP serves as a phosphate reserve under nutrient-limiting conditions (Plouviez *et al*., 2022). The second significant difference revealed by tomographic analyses concerned the electron-lucent vacuoles. Morphometric data showed that the vacuolar volume was increased by 7 to 9-fold in U200 relative to LoP or TAP (Fig. 2; Table S4). This result, attributable to U stress and not to phosphate limitation, was confirmed using the Cell Tracker Blue CMAC dye, which selectively labels vacuoles in plant cells (Stefano *et al*., 2018). The Cell Tracker Blue fluorescence signal in U200 was significantly higher than in LoP and revealed numerous punctuate foci attributable to vacuoles (Fig. S6). Enhanced vacuolization under U stress is consistent with a role of these organelles in stress mitigation. Increased vacuolar volume in microalgal cells exposed to high metal concentrations has been previously reported and attributed to either enhanced storage capacity for sequestering toxic ions or autophagy-mediated metal detoxification (Pérez-Pérez *et al*., 2017; Danouche *et al*., 2021). Autophagy is a membrane-trafficking process that delivers cytosolic components to the vacuole for degradation via autophagosomes. In green microalgae, this process is typically low under optimal conditions but is activated in response to a wide range of stress conditions, including metal stress (Pérez-Martín *et al*., 2015; Pérez-Pérez *et al*., 2017).

To gain insight into the ultrastructural changes induced by U, we analyzed its cellular distribution using TEM-EDX. No U could be detected in the EDX maps and spectra recorded for representative subcellular structures (cell wall, vacuole, acidocalcisome, chloroplast, mitochondrion, nucleus) of *Coelastrella* cells exposed to U200. Under these stress conditions, the cellular accumulation of U is transient since the alga can progressively release the metal after its rapid uptake from the medium (Beaulier *et al*., 2024). The relatively low sensitivity of TEM-EDX (Decelle *et al*., 2020), together with a low residual accumulation of U in *Coelastrella* cells after 24 h of stress could explain this negative result. To address this limitation, a more severe stress condition was used for chemical imaging of U. Cells were exposed to 400 µM uranyl nitrate (U400) in LoP for 24 h, a condition with no or limited U release by the alga, likely because its defense mechanisms are overwhelmed (Beaulier *et al*., 2024). As observed by TEM, exposure to U400 induced significant changes in cellular architecture relative to U200 (Fig. S5). The most striking changes were a stiffening of the cell wall (91 ± 10 nm for U200, 145 ± 24 nm for U400; n=4-5 cells) and the presence of multiple large vesicular structures generally located at the cell periphery (Fig. S5). A detailed analysis showed that this compartment contains diverse electron-dense cytoplasmic material sequestered in vacuolar structures, possibly linked to autophagy (Zhao *et al*., 2014; Couso *et al*., 2018). These ultrastructural changes were most likely a consequence of severe stress, in particular the accumulation of U resulting from limited detoxification capacity (Beaulier *et al*., 2024). EDX analysis of *Coelastrella* cells exposed to U400 allowed the detection of the characteristic M and L energy lines of U (around 3.17 and 13.61 keV, respectively) in three compartments (Fig. 3). Uranium was found in acidocalcisomes, and more specifically in the residual electron-dense matrix made of polyP granules, in the cell wall, and in autophagic-like vacuoles. In addition, extracellular precipitates containing U were observed in U400 samples (Fig. 3, S5). The formation of extracellular crystal structures containing U and potassium (K) has been described in the green microalga *Ankistrodesmus* sp. challenged with uranyl nitrate (Cheng *et al*., 2023). This observation was attributed to a possible biomineralization process, although the severe cellular damage induced by U limited further support for this interpretation. Careful examination of the EDX spectra recorded for *Coelastrella* cells challenged with U400 suggested that K may be associated with U in acidocalcisomes, the cell wall, autophagic-like vacuoles, and extracellular precipitates (Fig. 3). Because these precipitates lack crystalline structures, and discriminating the Kα and U-Mβ energy lines in EDX is challenging under the present conditions where K is not abundant, our data are not sufficient to conclude that U biomineralization occurs in *Coelastrella* sp. PCV, as suggested for *Ankistrodesmus* sp. (Cheng *et al*., 2023).

**Fig. 3:**
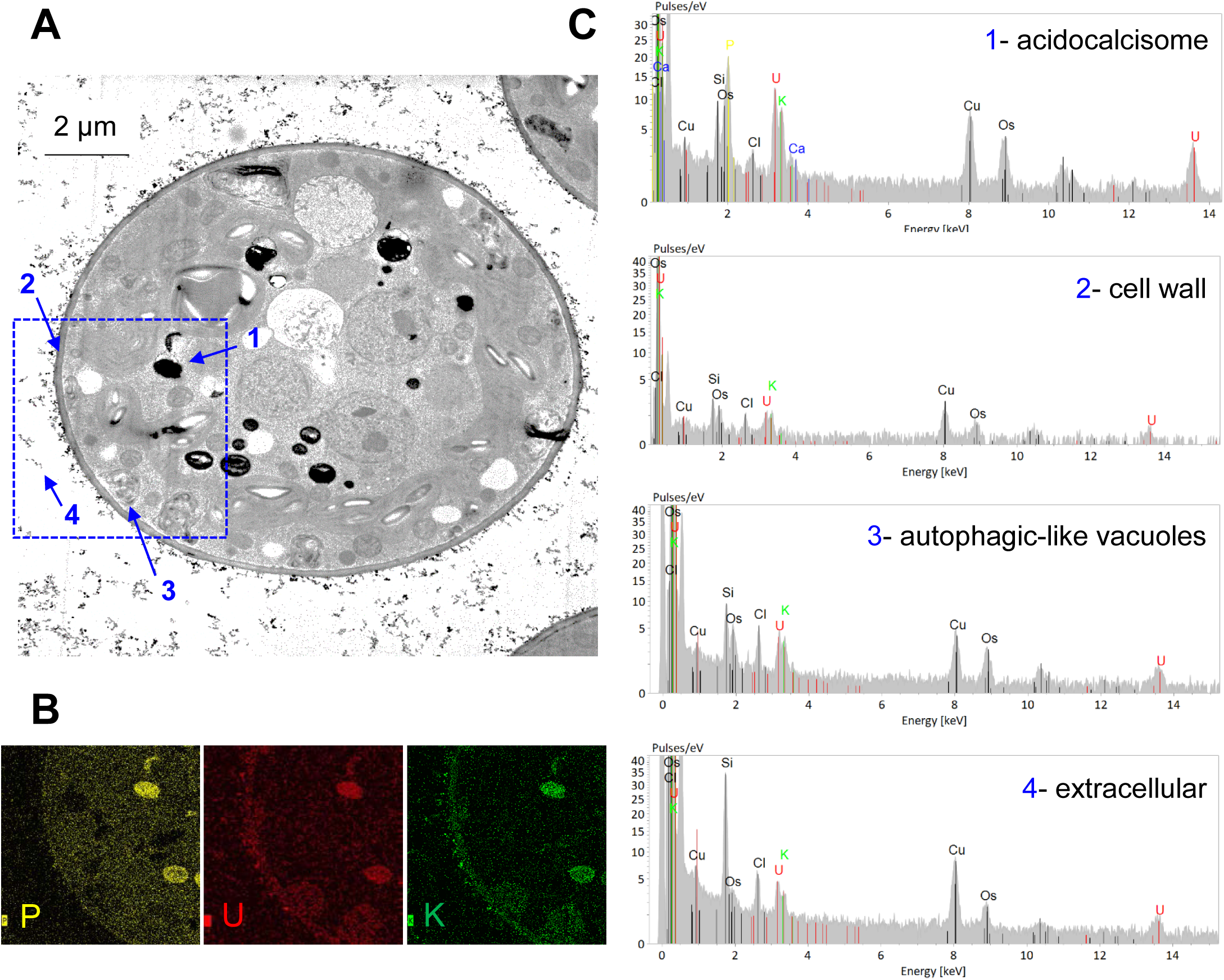
Structural and EDX analysis of *Coelastrella* exposed to U stress. Cells were grown for 24 h in U400, washed once with LoP before vitrification, and analyzed by TEM-EDX. **A** - TEM micrograph of a representative algal cell. Scale bar = 2 µm. **B** - EDX elemental maps of phosphorus, U and potassium in the region of interest (blue square). **C** - Representative EDX spectra obtained from the acidocalcisomes (1), cell wall (2), autophagic-like vacuoles (3), and extracellular precipitates (4). Elements indicated in black (e.g. osmium, copper) originate from the experimental setup (embedding resin or support grid).

### 3 Dynamics of the *Coelastrella* sp. PCV transcriptome under U stress

To investigate transcriptome dynamics under U stress, we established experimental conditions encompassing both the uptake and release phases of the toxic metal. In a typical experiment with 1 million cells/mL in LoP containing 200 µM uranyl nitrate, U accumulated very rapidly in cells, reaching a maximum at 1 h, followed by an efficient release into the medium resulting in low residual levels after 24 h (Fig. 4, S7). Under these conditions, both cell growth and photosynthesis were significantly inhibited by U (Fig. S8). Despite acute stress, algal vital functions were not irreversibly compromised, as evidenced by the near-complete recovery of photosynthesis after 48 h. For transcriptomic analysis, *Coelastrella* cells were harvested following 0.5, 1, 4 and 24 h of stress in U200. RNA libraries were prepared from four replicates of each sample and sequenced. A principal component analysis (PCA) of normalized gene expression values indicated that the first two components explained 63% of the variance in gene expression, with samples clustering along PCA1 according to the incubation time and along PCA2 according to the treatment (Fig. S9). Genes included in the differential expression analysis had a minimal expression of 1 FPKM in at least one time point in the experiment, exhibited a two-fold change (|log2 FC| ≥1) and a FDR ≤0.01 between control and treated samples at the same time point. A total of 5,289 genes were significantly regulated across the stress period (Table S5). The molecular response of *Coelastrella* to U200 was rapid, with 129 differentially expressed genes (DEGs) detected within 30 min, peaked at 4 h, and remained significant after 24 h of stress (Fig. S10). The number of up- and down-regulated genes were roughly similar at each time point. RNA-seq data were validated by qPCR for a subset of up- or down-regulated genes (Fig. S11).

**Fig. 4:**
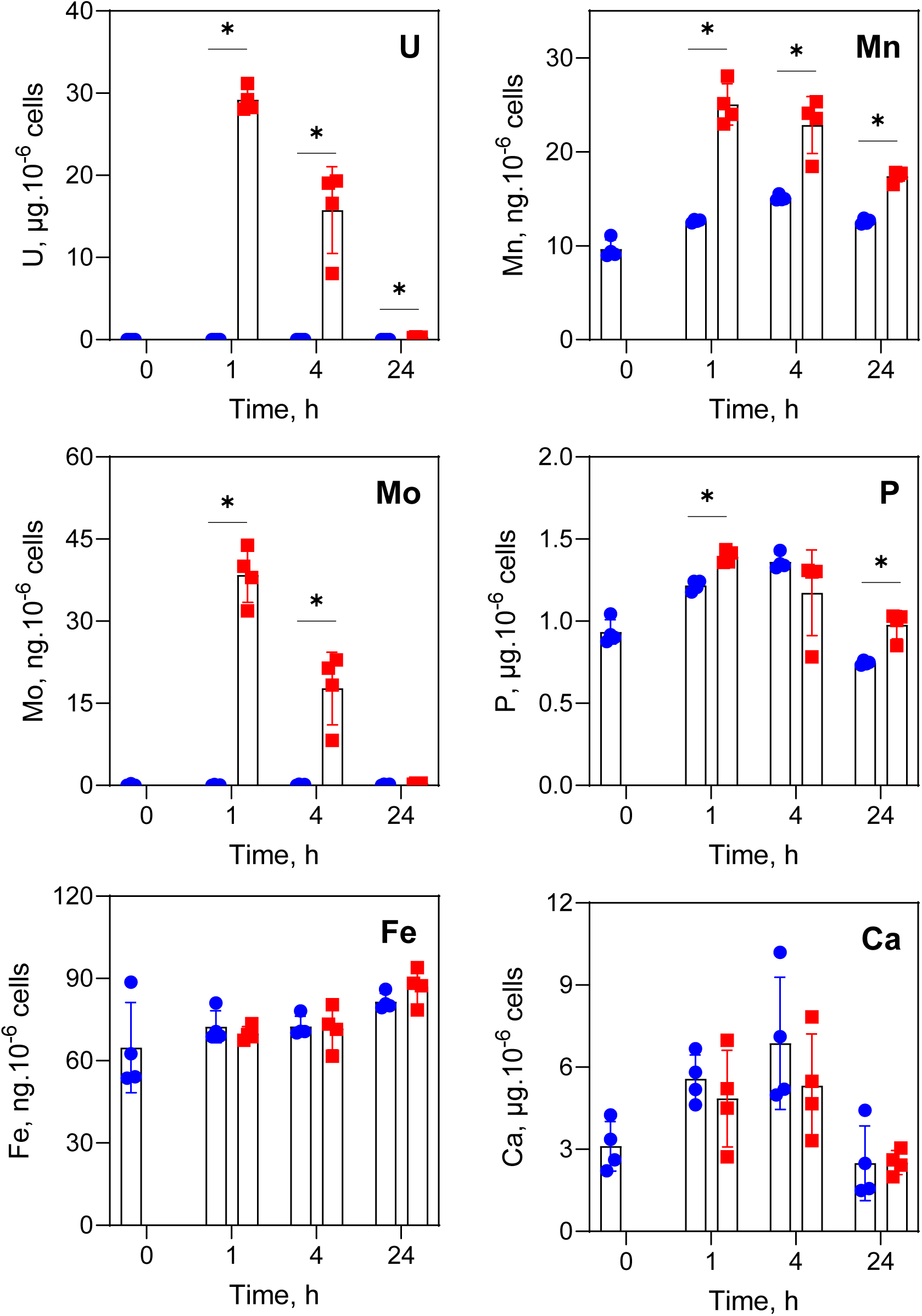
Ionomic analysis of *Coelastrella* cells challenged with U. Cells grown for 0, 1, 4, and 24 h in LoP (●) or U200 (▪) were harvested by centrifugation, washed with sodium carbonate, and mineralized in HNO_3_. Elemental composition was determined by ICP-MS and is expressed as µg or ng of element per million cells. Data were analyzed using the Mann-Whitney test. n=4 independent cultures per condition, identical to those used for RNA sequencing. Significance is indicated as p<0.05 (*).

To obtain a global view of transcriptome perturbations across biological processes, gene ontology (GO) term enrichment analysis was performed using BiNGO (Maere *et al*., 2005). Significantly enriched biological processes were identified for up- and down-regulated genes during the early (0.5 to 4 h) and late (24 h) phases of U stress response (Fig. 5). For clarity, only biological processes from the upper levels of the GO hierarchy are described. In the early phase of U stress, down-regulated genes were enriched in several GO terms related to protein synthesis (translation, GO:0006412; tRNA metabolic process, GO:0006399; ribosome biogenesis, GO:0042254) and cell division (cell cycle, GO:0007049; DNA metabolic process, GO:0006259; nucleoside monophosphate biosynthetic process, GO:0009124). During this phase, up-regulated genes were enriched in catabolism (catabolic process, GO:0009056; protein catabolic process, GO:0030163), transport (GO:0006810), in particular vesicle-mediated transport (GO:0016192), and lipid metabolic process (GO:0006629). The late phase of U stress was characterized by strong enrichment of GO terms related to photosynthesis (photosynthesis, GO:0015979; generation of precursor metabolites, GO:0006091; chlorophyll biosynthetic process, GO:0015995) for down-regulated genes, and to amino acid metabolism (GO:0006519) for up-regulated genes (Fig. 5).

**Fig. 5.**
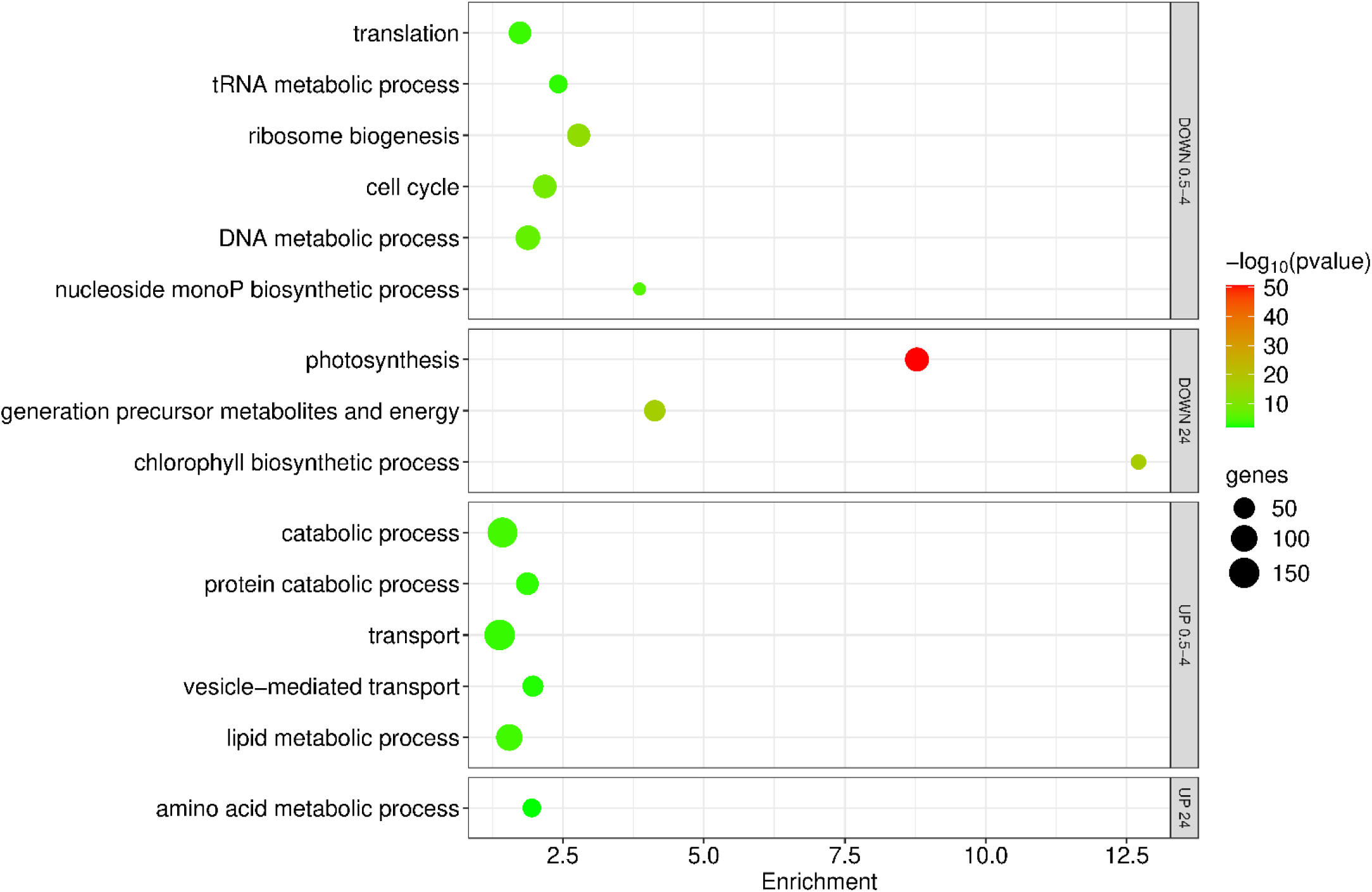
GO enrichment analysis of biological processes regulated by U stress in *Coelastrella*. GO enrichment analysis was performed using BiNGO with the following settings: statistical hypergeometric test, Bonferroni family-wise error rate multiple testing correction, and significant p-value <0.01. Bubble plots are separated in panels showing GO terms enriched for down- and upregulated genes in the early (0.5 to 4 h) and late (24 h) phases of U stress.

### 4 Molecular responses of *Coelastrella* sp. PCV to U stress

Here, we describe the molecular components linked to enriched GO terms and provide a detailed analysis of DEGs specifically associated with mineral homeostasis and metal stress. These biological processes are described through careful inspection and curation of the annotated *Coelastrella* sp. PCV genome, information from orthologous genes in the model organisms *C. reinhardtii* and *A. thaliana*, and functional data retrieved from various databases (e.g. UniProt, InterPro, TargetP-2.0).

#### Inhibition of cell division

The GO categories cell cycle (40 genes), microtubule-based movement (27 genes), DNA metabolic process (80 genes), and nucleoside monophosphate synthesis (20 genes) were enriched among down-regulated genes, indicating a significant effect of U on cell division (Fig. S12). This is supported by the inhibition of cell growth observed for *Coelastrella* exposed to U200 (Fig. S8) (Beaulier *et al*., 2024). Cell cycle related genes include four subunits of the condensing complex, two cyclins and five cyclin-dependent kinases, five components of the cohesin complex, and other cell cycle regulatory proteins (Fig. S12). Microtubule-based movement comprises 11 tubulin-related genes, 16 kinesin-like proteins, and five motor proteins dynein. The DNA metabolic process category was divided into DNA replication and DNA recombination, with repressed genes encoding many enzymes (e.g. helicases, DNA polymerase subunits) and regulatory factors. In addition, nine genes coding DNA repair proteins were down-regulated under U stress (Fig. S12). This indicates that exposure to U200 did not activate DNA repair pathways (e.g. correction of base mismatches or DNA lesions), suggesting that the stress was adequately managed by *Coelastrella* cells. As a comparison, a recent study showed that exposure of radish seedlings to moderate U doses (5-25 µM), thus excluding radiological effects, induced DNA damage and the up-regulation of genes associated with DNA repair in root meristem cells (Chen *et al*., 2025).

#### Imbalanced protein turnover and trafficking

Protein turnover emerged as a major cellular process influenced by U stress (Fig. 5). On the one hand, the non-overlapping GO categories translation (61 genes) and ribosome biogenesis (65 genes) were down-regulated under U200 stress, mainly at the end of the early stage (4 h). On the other hand, protein catabolic process (61 genes), vesicle-mediated transport (49 genes) and autophagy (10 genes) were up-regulated during the same period.

Down-regulated genes involved in translation encode for 24 proteins of the plastid ribosome subunits, eight eukaryotic translation initiation factor subunits, and 20 aminoacyl tRNA ligases, of which nine are predicted chloroplastic or mitochondrial enzymes (Fig. S13). These data suggest that chloroplast translation is preferentially inhibited under U stress, potentially placing chloroplast metabolism in a quiescent state while reducing the energy cost of protein synthesis. The ribosome biogenesis category comprises genes required for normal progression of rRNA processing, translation accuracy, and ribosome assembly, of which six genes annotated as ribosome biogenesis proteins and 15 genes involved in RNA modification, of which pseudouridine synthases and rRNA methyltransferases (Fig. S13).

Genes involved in protein catabolism and, in particular, in the ubiquitin-dependent protein catabolic process (46 genes) were significantly up-regulated in response to U stress (Fig. 5). For example, these genes encode eight E3 ubiquitin protein ligases, six E2 ubiquitin-conjugating enzymes, nine 26S proteasome non-ATPase regulatory subunits, and six proteasome alpha subunits (Fig. S13). The activation of ubiquitin-dependent protein degradation in response to U intoxication has been previously described in plants (Chen *et al*., 2025; Przybyla-Toscano *et al*., 2025).

Uranium stress in *Coelastrella* was also associated with a deregulation of vesicular-mediated protein transport (Fig. 5). Genes from this category were predominantly upregulated under U stress and encode vesicle trafficking proteins involved in the secretory pathway (Fig. S14). Some of these proteins are components of the ESCRT complex, which is primarily involved in the sorting of ubiquitinated plasma membrane proteins into multivesicular bodies for vacuolar degradation (Paez Valencia *et al*., 2016; Weiner *et al*., 2025). Other DEGs encode SNARE family and related proteins that are essential for the targeting and/or fusion of transport vesicles to their target membrane. Components of clathrin-dependent trafficking at plasma membrane, trans-Golgi network and endosomal system were also upregulated. Altogether, the upregulation of these pathways (Fig. S14) is consistent with enhanced sorting of membrane proteins for vacuolar degradation under U stress in *Coelastrella*. An important deregulation of the vesicular trafficking pathways has been recently described in Arabidopsis roots in response to U stress (Przybyla-Toscano *et al*., 2025), indicating that this response is conserved in plants and green microalgae. The ability of the ESCRT complex to bend membranes is also important for the maturation (closure) of autophagosomes during autophagy (Paez Valencia *et al*., 2016; Weiner *et al*., 2025). Consistent with this role, ten genes coding proteins forming the core autophagy machinery and participating in the formation of the autophagosome were up-regulated after 4 h of U stress (Fig. S14). As previously mentioned, this degradative process is activated under various stress conditions. For example, autophagy is triggered in *C. reinhardtii* challenged with silver (Pillai *et al*., 2014), copper, nickel or cobalt, but not with high concentrations of cadmium or mercury (Pérez-Martín *et al*., 2015). The activation of the autophagic pathway was also reported in root meristem cells from radish challenged with U (Chen *et al*., 2025).

#### Antioxidant response

Activation of the autophagic response under stress has been linked to the accumulation of reactive oxygen species (ROS) and redox imbalance (Pérez-Pérez *et al*., 2017). Uranium is known to promote the accumulation of ROS and to trigger oxidative stress in plants (e.g. (Vanhoudt *et al*., 2008; Vanhoudt *et al*., 2011; Tewari *et al*., 2015; Serre *et al*., 2019)). The antioxidant response to U is far less understood in microalgae (Pradines *et al*., 2005; Trenfield *et al*., 2012). Although the GO term ‘response to oxidative stress’ was not significantly enriched under U stress in *Coelastrella,* several genes coding antioxidant enzymes were identified among DEGs. This set included genes coding two chloroplastic iron(Fe)-dependent superoxide dismutases (SOD), one mitochondrial manganese(Mn)-dependent SOD, multiple chloroplastic peroxiredoxins, peptide methionine sulfoxide reductases and ascorbate peroxidases, three glutathione S-transferases (one chloroplastic), three glutathione peroxidases (one chloroplastic), two glutaredoxins (one chloroplastic), one monodehydroascorbate reductase, one catalase, and one sulfiredoxin (chloroplastic/mitochondrial) (Fig. S15). These genes displayed divergent expression patterns, either up- or downregulation, despite encoding proteins with identical functions. Immunoblot analysis revealed that the steady-state protein levels of Fe-dependent SOD and catalase did not change significantly under U stress, indicating that the transcriptional response was not directly reflected at the protein level for these two enzymes (Fig. S15). These results suggest that the antioxidant response triggered by U is complex, with probable transcriptional and post-transcriptional levels of regulation, time-dependent, gene-specific, and potentially compartment-specific, with the chloroplast emerging as a major site of the response. A similar situation was depicted in plants where severe U stress conditions induce changes in gene expression and enzyme capacities for different ROS-scavenging enzymes, including SOD, catalase and ascorbate peroxidase (Vanhoudt *et al*., 2008; Vanhoudt *et al*., 2011).

#### Metabolism

Nearly 700 genes related to metabolism were significantly affected by U stress. Enriched GO categories included amino acid metabolism (34 genes), lipid metabolism (101 genes), nucleoside synthesis (20 genes) and photosynthesis (73 genes) (Fig. 5). Because genes were both up and downregulated across diverse anabolic and catabolic pathways, and only one or a few DEGs were identified in multistep pathways, interpretation of transcriptomic data without information about changes in the corresponding metabolome remains mostly speculative and of limited insight. The impact of U on photosynthesis was the only effect clearly delineated, supported by physiological measurements, and is examined in detail. The photosynthetic electron transfer rate in *Coelastrella* cells challenged with U200 was significantly reduced after 24 h of stress, showing a 35% decrease compared to the control under light intensities exceeding 200 µEinstein (Fig. S8). At the transcriptome level, this inhibition was correlated with a widespread down-regulation of genes involved in photosynthesis light reactions (36 genes), generation of precursor metabolites (50 genes), and chlorophyll biosynthetic process (22 genes) (Fig. 5). All protein complexes of the thylakoid electron transfer chain were affected by gene down-regulation at 24 h (Fig. S16). Regulated genes are only nuclear-encoded. Immunoblot analysis showed a marked reduction in PsaD steady-state level after 24 h of stress (Fig. S16), corroborating the transcriptional down-regulation of the corresponding gene and aligning with *in vivo* photosynthesis measurements (Fig. S8). In addition, carbon fixation was affected, with downregulation of genes coding nearly all enzymes of the Calvin cycle and the regulatory protein CP12 after 24 h of U stress (Fig. S16). Also, chlorophyll biosynthetic and catabolic processes were significantly impacted by U stress. On the one hand, 17 genes involved in chlorophyll biosynthesis were downregulated (Fig. S17). Notably, the *FLU* gene was rapidly downregulated, with repression observed as early as 1 h, and maintained at 4 and 24 h of stress. In *Chlamydomonas*, the FLU1-like protein regulates 5-aminolevulinic acid synthesis, the universal precursor of all tetrapyrroles (Falciatore *et al*., 2005). On the other hand, nine genes coding pheophorbide a oxygenase, which catalyzes the key reaction of chlorophyll catabolism, were upregulated (Fig. S17).

#### Mineral homeostasis, metal transporters and metalloproteins

To gain insight into the effects of U on mineral homeostasis, we combined transcriptomic and ionomic analysis of *Coelastrella* cells under U stress. Ionomic analysis was performed on the same samples used for RNA-seq to ensure accurate integration of response dynamics (Fig. 4) and was replicated in independent experiments to ensure reliable interpretation. Samples for ICP-MS analysis consisted of whole cells washed with sodium carbonate to remove loosely bound U (Beaulier *et al*., 2024), reflecting the total cellular fraction of elements, including both cell wall-bound and internalized pools.

### Manganese

The Mn content in *Coelastrella* cells increased rapidly and remained elevated under U stress, showing a 2-fold increase at 1 h and about 1.5-fold at 4 and 24 h (Fig. 4). Despite this, there was a limited effect of U on genes related to Mn homeostasis, with differential expression observed in two genes coding metal transporters and six genes coding metalloproteins (Fig. 6A). Two genes orthologous to the *C. reinhardtii CMT1A* and *PAM71* genes, respectively implicated in Mn uptake into chloroplasts and Mn transport across thylakoid membranes (Schneider *et al*., 2016), were repressed after 4 to 24 h of stress. No transporter that could be involved in Mn uptake from the medium was identified among DEGs. The six genes coding Mn-binding proteins were either repressed (1-deoxy-D-xylulose 5-phosphate reducto-isomerase, protein phosphatase 2C, photosystem II core complex protein PsbY1) or induced (two leucine aminopeptidases; mitochondrial Mn-SOD) (Fig. 6A). Although the subcellular distribution of Mn in *Coelastrella* cells is not known, these data suggest that Mn accumulated during U stress may be preferentially allocated to sustain Mn-dependent SOD activity in mitochondria and maintain redox homeostasis (Allen *et al*., 2007). This may reduce Mn allocation to chloroplasts, where the demand for building the MnCaOcluster of the oxygen-evolving complex is likely reduced under U stress (Fig. S8, S16). In plants, moderate to high U stress triggers substantial perturbations in the homeostasis of many mineral elements. Under these conditions, the Mn content is either not affected or decreased in roots, and eventually in leaves (Vanhoudt *et al*., 2011a; Lai *et al*., 2020). This response parallels that of several other elements, suggesting a lack of specificity, and contrasts with the increased levels described in *Coelastrella* cells (Fig. 4).

**Fig. 6:**
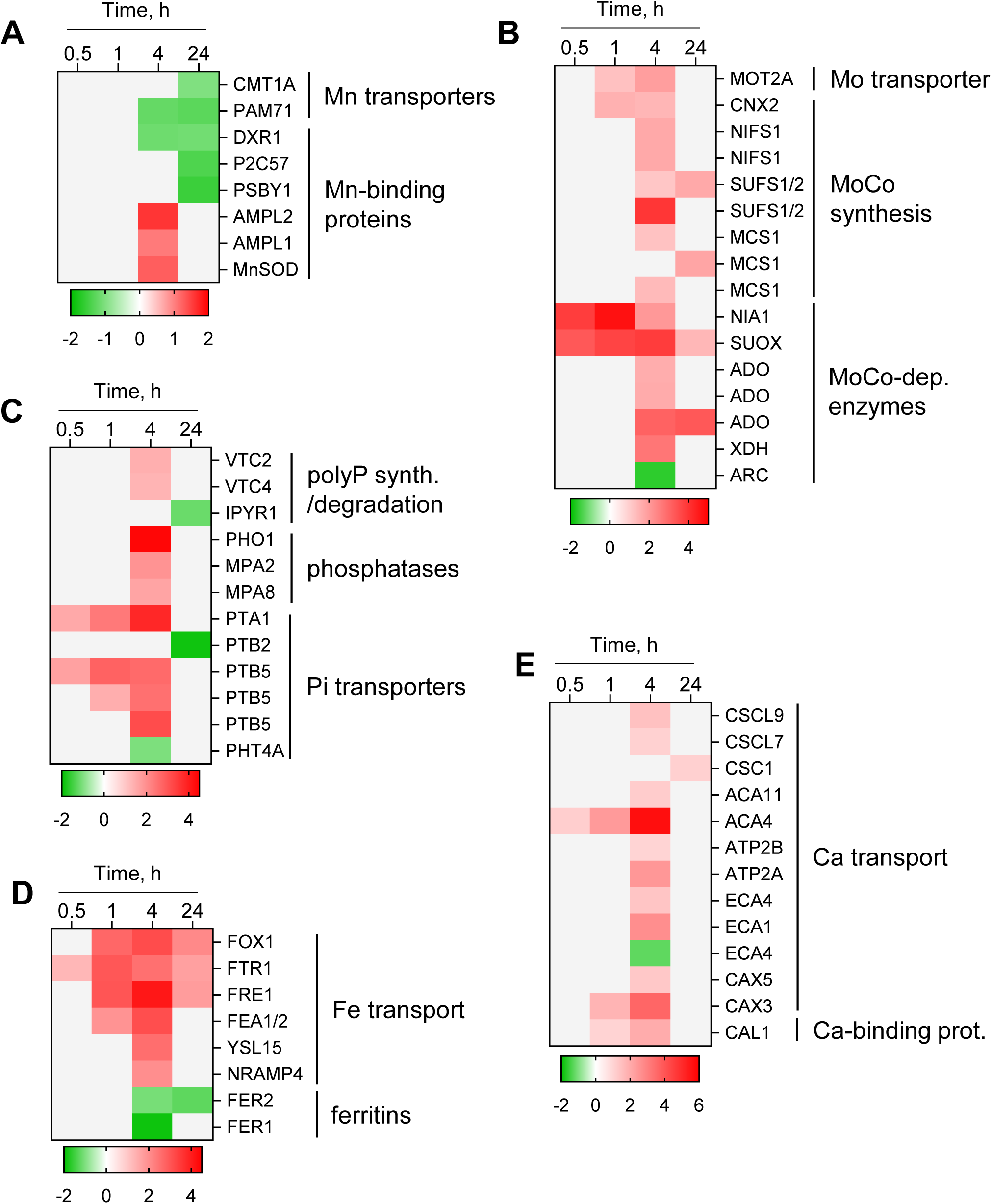
Regulation of genes involved in mineral homeostasis. Heatmaps are illustrating the expression of genes involved in Mn (**A**), Mo (**B**), Pi (**C**), Fe (**D**), and Ca (**E**) homeostasis. Genes showing significant changes in expression in response to U stress (FPKM ≥1, |log_2_FC| ≥1, FDR ≤0.01) are colored red (upregulated) or green (downregulated). Non-significant changes are shown in light grey. Expression values are displayed on a log_2_ scale.

### Molybdenum

The homeostasis of molybdenum (Mo) was markedly modified in *Coelastrella* cells challenged with U. The accumulation of Mo closely paralleled that of U, showing a >300-fold increase at 1 h, a 150-fold increase at 4 h, and a return to control levels at 24 h (Fig. 4). Such perturbations of Mo homeostasis in response to U stress remain poorly documented (Sarthou *et al*., 2020; Han *et al*., 2025). Transcriptomics revealed the molecular basis of Mo changes in *Coelastrella*. First, a high affinity molybdate transporter coding gene (*MOT2A*) was upregulated after 1 and 4 h of stress, suggesting an improved uptake capacity in *Coelastrella* cells (Fig. 6B). This hypothesis is supported by an increased withdrawal of Mo from the medium by U-treated cells compared to control cells (Fig. S7). Second, genes coding several enzymes involved in the biosynthesis of the molybdenum cofactor (MoCo), the biologically active form of this transition metal (Kaufholdt *et al*., 2017), were upregulated at 4 and 24 h. These genes encode a GTP 3’,8-cyclase (CNX2) catalyzing the first step in mitochondria, four potential cysteine desulfurases that provide the sulfur atom for MoCo biosynthesis (NIFS1, SUFS1/2), and three MoCo sulfurases (MCS1) (Fig. 6B). Third, genes coding enzymes from the two families of MoCo-dependent proteins found in the green linage (Kaufholdt *et al*., 2017; Tejada-Jimenez *et al*., 2023) were markedly induced as early as 30 min after the onset of U stress. They encode two enzymes of the sulfite oxidase family, a nitrate reductase (NR, *NIA1* gene) and a sulfite oxidase (SUOX), and four enzymes of the xanthine oxidoreductase family, three probable aldehyde dehydrogenases/oxidases (ADO) and a xanthine dehydrogenase (XDH). The only down-regulated gene related to Mo homeostasis encodes the amidoxime-reducing component (ARC) (Fig. 6B), a molybdoenzyme catalyzing the reduction of various compounds (Tejada-Jimenez *et al*., 2023). Taken together, these findings suggest a substantial demand for MoCo starting very rapidly after the beginning of U stress. This demand is likely to sustain the first step of nitrate assimilation (NR), the detoxification of sulfite and the potential regulation of the balance between sulfur and nitrogen metabolism (SUOX), the catabolism of purines (XDH), and possibly the detoxification of carbonyl aldehydes (ADO) (Tejada-Jimenez *et al*., 2023). In *Chlamydomonas*, NR and ARC work together in the cytoplasm for the reduction of nitrite into nitric oxide, and the corresponding genes are coordinately expressed (Chamizo-Ampudia *et al*., 2016). Nitric oxide has been implicated in many important cellular processes in photosynthetic organisms, of which acclimation to abiotic stresses, and accumulates in Arabidopsis roots challenged with U (Tewari *et al*., 2015; Serre *et al*., 2019). In *Coelastrella* exposed to U200, the opposite regulation of *NR* (induction) and *ARC* (repression) genes precludes firm conclusions about the involvement of nitrite-dependent nitric oxide production in metal stress response.

### Phosphate

Interactions between U and inorganic phosphate (Pi) are known as a critical factor influencing U speciation, bioavailability and toxicity. As a consequence, the control medium used in this study was a modified TAP containing low Pi, and DEGs uncovered by the RNA-seq analysis highlight transcriptional changes triggered by U200 under Pi-limiting conditions. The amount of phosphorus measured in whole *Coelastrella* cells remained relatively stable throughout the experimental period and was only marginally affected by U stress (Fig. 4). These data are consistent with the similar morphometric characteristics of polyP granules in LoP and U200 (Fig. 2, S4, Table S3), suggesting that the mobilization of Pi reserves to cope with Pi limitation is comparable in both conditions. To support this view, only three genes potentially involved in polyP synthesis (*VTC2* and *VTC4*) or degradation (IPYR1, candidate soluble inorganic pyrophosphatase 1) (Plouviez *et al*., 2023) were moderately deregulated in the presence of U (Fig. 6C). In contrast, genes coding two acidic phosphatases (MPA2 and MPA8) and one alkaline phosphatase (PHO1) were highly induced after 4 h of stress. This suggests a compensatory mechanism for Pi acquisition from organic sources. In support to this hypothesis, it was shown that U triggers a depletion of both Pi and phosphorylated metabolites in *Arabidopsis* cells (Berthet *et al*., 2018). In addition, U significantly altered the expression of genes coding Pi transporters. Orthologs of the *C. reinhardtii* genes coding a sodium/Pi symporter (*PTB5*) and a proton/Pi symporter (*PTA1*) were upregulated, whereas orthologs of *PTB2* and *PHT4A* were downregulated (Fig. 6C). The probable localization of these transporters at the plasma membrane, together with their transcriptional regulation in response to Pi availability (Sanz-Luque & Grossman, 2023), suggests that the intracellular Pi depletion was perceived as more severe in U200 than LoP and that *Coelastrella* cells attempted to increase their capacity for external Pi uptake under U stress. This assumption is supported by the faster withdrawal of Pi from the medium in U-treated cells relative to control cells (Fig. S7).

### Iron

Several genes responding to Fe deficiency in green microalgae or land plants were regulated in *Coelastrella* exposed to U. Orthologous genes of the *C. reinhardtii* reduction-oxidation transport mechanism (Liu *et al*., 2024) consisting of an Fe(III)-chelate reductase/oxidoreductase (FRE1), an Fe(II) reductase (FOX1), and an Fe(III) permease (FTR1) were induced very rapidly after stress initiation (Fig. 6D). A similar trend was observed for a potential extracellular Fe-assimilation protein (FEA1/2) (Allen *et al*., 2007) and an oligopeptide transporter (OPT) orthologous to Yellow Stripe-like Fe-siderophore transporters from *Oryza sativa* (YSL15) (Inoue *et al*., 2009). The upregulation of these genes is consistent with a cellular response aimed at enhancing Fe uptake. Another hypothetical Fe transporter-coding gene, upregulated by 4-fold after 4 h of U stress, is orthologous to the *C. reinhardtii* NRAMP4 protein that may be involved in the remobilization of vacuolar Fe stores in *C. reinhardtii* (Urzica *et al*., 2012). In addition, many genes coding Fe-containing proteins were either upregulated (51 genes) or downregulated (36 genes), suggesting an important perturbation of Fe homeostasis by U. Among them, FER1 and FER2 genes coding ferritins were down-regulated under U stress (Fig. 6D). This response is the opposite to the transcriptional signature of Fe deficiency in *C. reinhardtii* (Urzica *et al*., 2012) but similar to that described in marine species *Dunaliella*, in which FER1 transcript and protein are down-regulated in response to Fe deprivation (Davidi *et al*., 2023).

Despite the significant perturbation of Fe homeostasis genes, ionomic analysis showed no significant changes in total Fe content in *Coelastrella* cells or in Fe withdrawal from the medium in U200 compared to LoP (Fig. 4, S7). As a consequence, one can hypothesize a redistribution of Fe between subcellular compartments, resulting in the sensing of Fe deficiency and the transcriptional regulation of key Fe-responsive genes. The capacity of acidocalcisomes to accumulate U (Fig. 3) and store Fe (Schmollinger *et al*., 2021) may suggest that Fe from acidocalcisomes could be reallocated to subcellular compartments with higher Fe demands. NRAMP4 might be involved in the mobilization of Fe from acidocalcisomes. Such a scenario has been hypothesized in *Arabidopsis* challenged with U, involving a redistribution of Fe from the vacuole to chloroplasts (Berthet *et al*., 2018). However, the situations described in *Coelastrella* and *Arabidopsis* result in opposite responses of the Fe-regulated networks, as U stress is perceived as an excess of Fe in Arabidopsis, leading to the downregulation of Fe-responsive genes (Doustaly *et al*., 2014). The induction of the Fe(III) uptake machinery during U stress in *Coelastrella* is paradoxical and raises questions about how cells manage the uptake of the toxic element. Indeed, the high affinity Fe(III) permease Ftr1 has been implicated in U uptake in *Saccharomyces cerevisiae* (Revel *et al*., 2022), and if this mechanism is conserved in green microalgae, the upregulation of *FTR1* would indicate an increased capacity of *Coelastrella* to take up the radionuclide. Also, the Fe(III) transporter YSL15 has been involved in mediating chromium(III) uptake in rice (Li *et al*., 2024), suggesting that YSL15 from *Coelastrella* could contribute to uranyl uptake, as both Fe(III), chromium(III) and UO_2_^2+^ are all classified as hard Lewis acids. If FTR1 and/or YSL15 contribute to uranyl uptake in *Coelastrella*, the accumulation of the metal in cells is however transient (Fig. 4), indicating that the uptake is rapidly counteracted by a highly efficient efflux process. Candidates transporters involved in this efflux are discussed below.

### *Calcium

The interaction between U and Ca is well established in higher plants, in which U uptake is mediated by Ca-permeable cation channels and U distribution in tissues is Ca dependent (El Hayek *et al*., 2019; Rajabi *et al*., 2021; Mertens *et al*., 2022; Sarthou *et al*., 2022). In *C. reinhardtii*, Ca was shown to inhibit uranyl uptake, suggesting that Ca assimilation routes could serve for U absorption (Fortin *et al*., 2007). In *Coelastrella*, U stress triggered the deregulation of 50 genes related to Ca (21 downregulated, 29 upregulated), suggesting a disruption of its homeostasis and associated cellular processes. First, genes coding putative osmo-sensitive (*CSCL7*, *CSCL9*) and stress-gated (*CSC1*) Ca-permeable channels were moderately upregulated after 4 or 24 h of stress (Fig. 6E). Because the total amount of Ca in *Coelastrella* cells was not affected by U stress (Fig. 4), transcriptional deregulation of these genes is expected to exert limited effects on Ca uptake from the medium. Second, nine genes coding transporters potentially involved in maintaining cytosolic Ca concentrations were deregulated, mostly upregulated, under U stress. Seven genes are coding putative Ca-transporting P-type ATPases, including four P-IIA endoplasmic reticulum (ER)-type Ca^2+^ ATPases (ECA) and three PII-B autoinhibited Ca^2+^ ATPases (ACA). Although the subcellular localization of these transporters in not known in *C. reinhardtii*, they are predicted to catalyze the translocation of Ca from the cytosol to the ER lumen, the organelles (vacuole, chloroplast, mitochondrion) or out of the cell (Pivato & Ballottari, 2021). All but one of these genes were upregulated during U stress (Fig. 6E), of which an ortholog of the *Arabidopsis ACA4* gene that is involved in the translocation of Ca from the cytosol to small vacuoles (Geisler *et al*., 2000). The expression of the *Coelastrella ACA4* gene was rapidly induced, starting 30 min after the onset of U stress, and strongly upregulated, reaching a 50-fold increase at 4 h relative to the control. In addition to P-type ATPases, genes orthologous to the Ca/proton exchanger *CAX3* and the Ca/sodium exchanger *CAX5* from *C. reinhardtii* were upregulated in response to U (Fig. 6E). *CAX3* displayed the earliest and strongest response to U, and the encoded protein was predicted to localize to the endomembrane system. Transcriptional regulation of genes coding both Ca-transporting ATPases and Ca exchangers likely contributes to the maintenance of Ca homeostasis during U stress, as previously described in Arabidopsis (Mertens *et al*., 2022). Also, since Ca channels can transport U (Revel *et al*., 2022; Sarthou *et al*., 2022), one can hypothesize that some of these transporters, notably those showing the strongest upregulation during U stress (ACA4 and CAX3), may contribute to the translocation of U from the cytosol to the acidocalcisomes or vacuoles (Fig. 3) or to its export outside algal cells (Fig. 4). Last, U stress also affected genes coding Ca-binding proteins as well as Ca- and/or calmodulin-regulated proteins, including protein kinases. Among them, the CAL1 protein harboring a putative Ca-binding calreticulin/calnexin domain at the C-terminus and a catalytic domain of calmodulin-dependent serine/threonine kinases at the N-terminus was upregulated (Fig. 6E). Given that calreticulin is a high-capacity Ca-binding protein and the uranyl cation can compete with Ca for binding to specific proteins (Garai & Delangle, 2020; Vallet *et al*., 2023), one can postulate that CAL1 from *Coelastrella* may interact with U.

#### Additional transporters potentially involved in U detoxification

In plants and algae, ATP-binding cassette (ABC) and multidrug and toxic compound extrusion (MATE) transporters play key roles in metal homeostasis and heavy metal detoxification by mediating the translocation of ions and their conjugates across membranes (Upadhyay *et al*., 2019; Do *et al*., 2021; Li *et al*., 2022). The expression of 30 *Coelastrella* genes annotated as ABC transporters was affected by U, with 27 being upregulated. These genes were further assigned to seven subclasses according to (Li *et al*., 2022). Although ABC transporters exhibit broad and often pleiotropic transport activities, this classification guides predictions of potential substrates and roles in heavy metal stress responses. Accordingly, only a few genes from subclasses C and G are proposed to participate to U detoxification (Fig. 7). The three upregulated C-class genes may contribute to U transport, as they operate as xenobiotic-transporting ATPases. Indeed, the *Coelastrella ABCC3/4/8* genes are orthologs of the *A. thaliana ABCC1/2* genes that contribute to detoxification by sequestering heavy metals such as arsenic, cadmium and mercury in vacuoles (Park *et al*., 2012). Ten *ABCG* genes, also known as pleiotropic drug resistance (PDR) transporters, were also upregulated. In plants, PDR-type ABCGs participate in cell detoxification, metabolite excretion, and responses to biotic and abiotic stresses (Do *et al*., 2021). Several of these genes were strongly induced (> 12-fold) in U200 conditions, notably *ABCG23* at 4 h and *ABCG16* at 24 h (Fig. 7), consistent with a potential role in the secretion of U or U-chelating metabolites into acidocalcisomes or the extracellular medium. Finally, four genes encoding MATE transporters were induced between 4 and 24 h of U200 stress (Fig. 7). Given the broad substrate specificity of these transporters, it is difficult to assess their specific role in the detoxification of U. However, the most strongly upregulated gene (*DTX42*) is orthologous to the root citrate transporters MATE and FRD3 involved in aluminum tolerance (Liu *et al*., 2009) and Fe loading into the xylem (Durrett *et al*., 2007), respectively. This suggests that, in *Coelastrella,* DTX42 may facilitate the efflux of an organic acid to chelate uranyl ions in the extracellular environment, contributing to the mitigation of U stress.

**Fig. 7:**
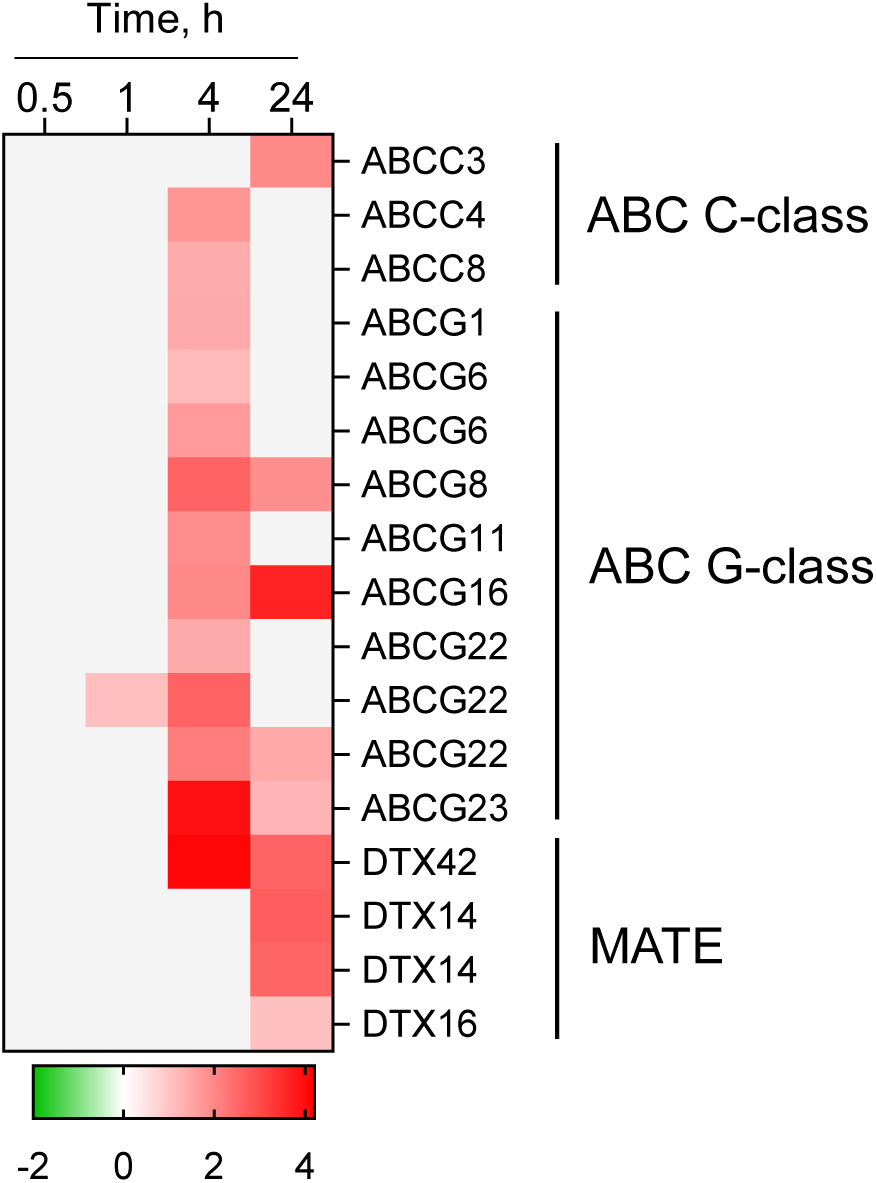
Regulation of genes coding ABC and MATE transporters potentially involved in U detoxication. Genes showing significant changes in expression in response to U stress (FPKM ≥1, |log_2_FC| ≥1, FDR ≤0.01) are colored red (upregulated) or green (downregulated). Non-significant changes are shown in light grey. Expression values are displayed on a log_2_ scale.

## Conclusion

By integrating high-resolution imaging with multi-omics analyses, this study provides a comprehensive understanding of how the metal-tolerant microalga *Coelastrella* sp. PCV responds to U chemotoxicity. Our analyses delineate the cellular and molecular consequences of U intoxication while uncovering the cellular fate of U in *Coelastrella* and the molecular mechanisms that may mediate its detoxification (Fig. 8). Growth inhibition and photosynthetic decay result from the massive repression of genes involved in cell cycle regulation, DNA metabolic process, and photosynthesis (Fig. 5). Although the inhibition of photosynthesis by U has been reported in microalgae (Herlory *et al*., 2013) and plants (Gao *et al*., 2019), the underlying molecular mechanisms have remained unknown. Here, we show that photosynthetic impairment reflects the coordinated repression of nuclear genes for thylakoid electron transfer and stromal carbon fixation, together with a reduced capacity for chlorophyll accumulation (Fig. S16, S17). The downregulation of ribosome biogenesis and translation, along with the upregulation of ubiquitin-related pathways, vesicle-mediated transport and autophagy, supports the view that U stress also induces a major imbalance in protein turnover and trafficking. The induction of autophagy-related genes (Fig. S14), the enhanced vacuolization (Fig. 2, S6), and the detection of U in autophagic vacuoles (Fig. 3) collectively demonstrate that U stress in *Coelastrella* triggers significant sorting of proteins for vacuolar degradation.

**Fig. 8:**
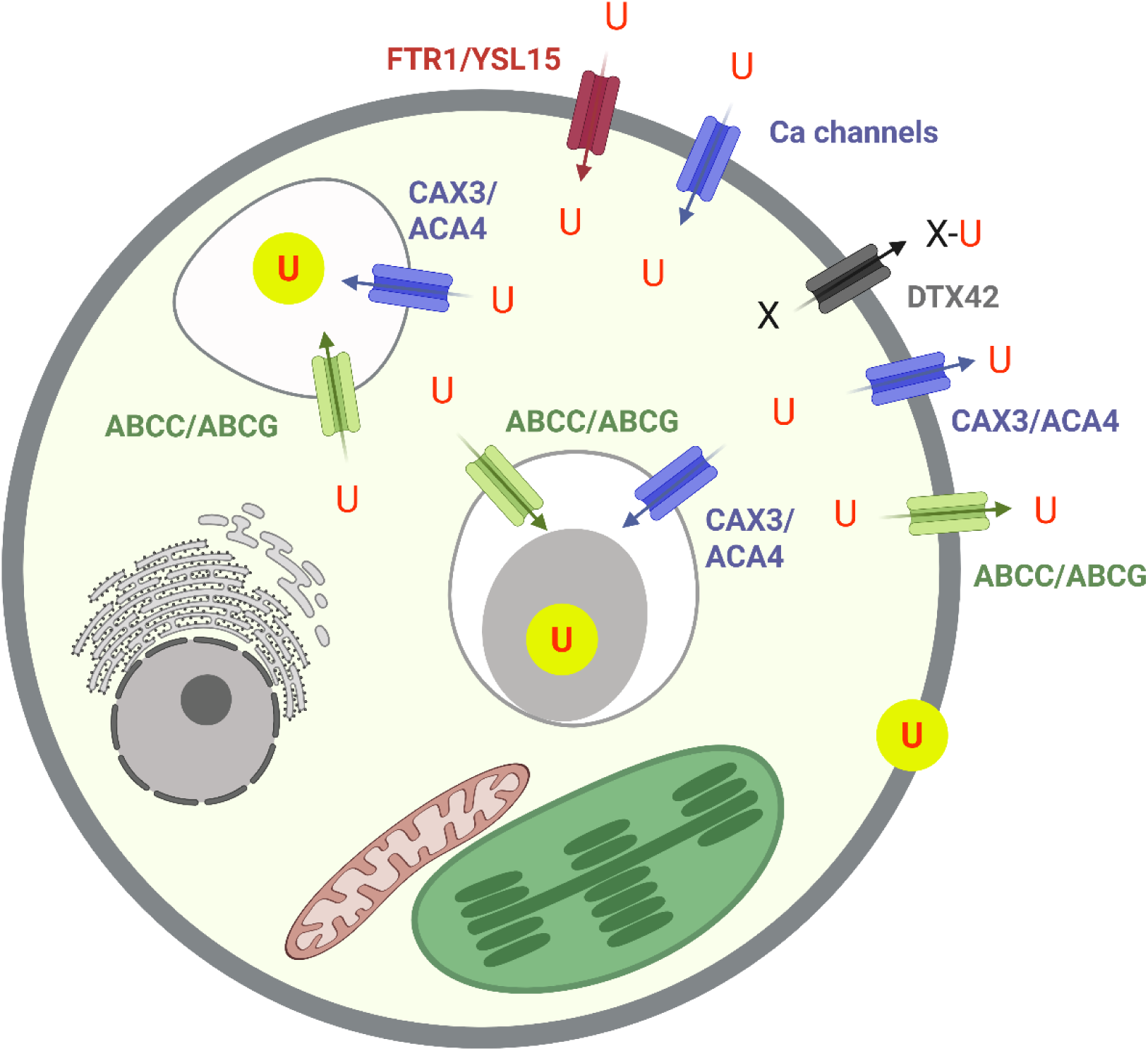
Schematic representation of the proposed molecular mechanisms involved in U tolerance in *Coelastrella* sp. PCV. Uranium was localized to the cell wall, polyphosphate bodies within acidocalcisomes, and vacuoles. Transporters potentially involved in U uptake, intracellular trafficking and efflux were identified among strongly deregulated genes (Fig. 6, 7). Transporters associated with Ca and Fe homeostasis are shown in blue and red, respectively. Members of the ABC and MATE transporter families are in green and dark grey, respectively.

Transcriptomic and ionomic analyses also shed light on the consequences of U intoxication on the homeostasis of essential elements (Fig. 4, 6). The strong and rapid induction of Mo uptake, MoCo synthesis, and MoCo-utilizing enzymes has never been reported in any organism exposed to U. Although MoCo-dependent reactions play central metabolic roles (e.g. nitrate assimilation, detoxification of sulfite and carbonyl aldehydes), interpretation of these changes in Mo homeostasis remains unclear. In addition, U markedly rewired gene networks controlling the homeostasis of Mn, Pi, Fe or Ca. While some responses mirror those previously reported in plants (e.g. (Doustaly *et al*., 2014; Lai *et al*., 2020)) and bacteria (Kolhe *et al*., 2018), these regulatory patterns exhibit unique specificities. Structural remodeling of polyP bodies in acidocalcisomes, together with the induction of genes coding phosphatases and plasma membrane Pi transporters, highlights compensatory mechanisms for Pi acquisition from organic sources and from the external environment under U stress. Genes contributing to Fe uptake and intracellular trafficking were also strongly upregulated, consistent with a Fe-starvation-like response, resulting in a probable redistribution of Fe among subcellular compartments. Similarly, deregulation of Ca-transporting ATPases and Ca exchangers is likely required to maintain Ca homeostasis across distinct cellular structures. Beyond this role, Ca transporters may also facilitate U sequestration into acidocalcisomes (Fig. 8). Indeed, our findings indicate that these structures, and more specifically their matrix made of polyP bodies, not only maintain essential ion homeostasis (Hong-Hermesdorf *et al*., 2014; Tsednee *et al*., 2019; Schmollinger *et al*., 2021) but also modulate the intracellular concentration of hazardous metals via compartmentalization and sequestration. Last, a striking feature of *Coelastrella* is its ability to rapidly and efficiently release U upon exposure to concentrations that are lethal to non-tolerant microalgae (Beaulier *et al*., 2024). The efflux of U may involve Ca transporters and some members of the ABC and MATE transporters that are upregulated in response to the metal (Fig. 8).

The recent development of a transformation method for a *Coelastrella* species (Baldanta *et al*., 2025) paves the way for the functional characterization of candidate genes and proteins identified in this study. Elucidating tolerance and accumulation mechanisms in *Coelastrella* sp. PCV could, in turn, drive innovations in green biotechnology, including bioremediation of metal-contaminated environments, sustainable nanoparticle biosynthesis (Choudhary *et al*., 2023), and metal capture for biomedical or industrial applications using engineered metal-binding proteins integrated into specialized devices.

## Acknowledgements

This work was funded by the Agence Nationale de la Recherche (ANR-21-CE34-0004, DemoniaCo, and ANR-17-EURE-0003, CBH-EUR-GS), the INRAE Plant Biology and Breeding division, and the CNRS Biologie program “Diversity of biological mechanisms”. CB was supported by a PhD fellowship from the IDEX University Grenoble Alpes.

We thank Gaelle Villain (LPCV, Grenoble, France) for the maintenance of the algal growth facility and Benoit Gallet (IBS, Grenoble, France) for assistance in the cryo-preparation of cell samples. We acknowledge the team at Sequentia Biotech for their sequencing work, data-mining support, constructive discussions, and consistent availability throughout this project.

## Author contributions

CB performed the experiments and contributed to data analysis and interpretation. FD, AG, GSL and PHJ performed the experiments and contributed to data analysis. JPT and CA contributed to data interpretation and manuscript editing. SR designed the research, contributed to data interpretation, and wrote the manuscript.

## Supporting information

**Table S1**: Genomic datasets used in this study.

**Table S2**: Genome assembly evaluation.

**Table S3**: Primers used for QPCR analysis.

**Table S4:** Morphometric analysis of subcellular structures in Coelastrella exposed to U.

**Table S5:** Differentially expressed genes in *Coelastrella* sp. PCV exposed to U200.

**Fig. S1:** Assessment of the *Coelastrella* sp. PCV genome completeness using BUSCO.

**Fig. S2:** Structure and annotation of the assembled *Coelastrella* sp. PCV chloroplast (A) and mitochondrial (B) genomes.

**Fig. S3:** EDX analysis of acidocalcisomes in *Coelastrella* sp. PCV.

**Fig. S4:** Morphometric parameters of acidocalcisomes and polyP granules in *Coelastrella* cells challenged with U.

**Fig. S5:** TEM micrographs of *Coelastrella* in LoP, U200 and U400 conditions.

**Fig. S6:** Observation of vacuoles in *Coelastrella* cells using Cell Tracker Blue.

**Fig. S7:** Elemental composition of growth media used for *Coelastrella* cells exposed to U.

**Fig. S8:** Growth and photosynthesis of *Coelastrella* cells challenged with U.

**Fig. S9:** PCA of normalized gene expression values for all samples.

**Fig. S10:** Dynamics of differential gene expression during the U stress response in *Coelastrella*.

**Fig. S11:** QPCR analysis of DEGs identified through RNA-seq.

**Fig. S12:** Regulation of genes involved in cell division.

**Fig. S13:** Regulation of genes involved in protein turnover.

**Fig. S14:** Regulation of genes involved in vesicle-mediated transport (A) and autophagy (B).

**Fig. S15:** Regulation of genes and proteins involved in the antioxidant response.

**Fig. S16:** Regulation of genes and proteins involved in photosynthesis.

**Fig. S17:** Regulation of genes involved in chlorophyll synthesis (A) and catabolism (B).

**Movie S1:** 3D reconstructed *Coelastrella* sp. PCV cell grown in mixotrophic conditions.

