## Supplemental Tables and figures for "Integrated cellular and molecular responses to uranium chemotoxicity in the metal-tolerant microalga *Coelastrella* sp. PCV"

***New Phytologist* Supporting Information**

The following Supporting Information is available for this article:

**Table S1**: Genomic datasets used in this study.

**Table S2**: Genome assembly evaluation.

**Table S3**: Primers used for QPCR analysis.

**Table S4:** Morphometric analysis of subcellular structures in Coelastrella exposed to U. *Excel file.*

**Table S5:** Differentially expressed genes in *Coelastrella* sp. PCV exposed to U200. *Excel file.*

**Fig. S1:** Assessment of the *Coelastrella* sp. PCV genome completeness using BUSCO.

**Fig. S2:** Structure and annotation of the assembled *Coelastrella* sp. PCV chloroplast (A) and mitochondrial (B) genomes.

**Fig. S3:** EDX analysis of acidocalcisomes in *Coelastrella* sp. PCV.

**Fig. S4:** Morphometric parameters of acidocalcisomes and polyP granules in *Coelastrella* cells challenged with U.

**Fig. S5:** TEM micrographs of *Coelastrella* in LoP, U200 and U400 conditions.

**Fig. S6:** Observation of vacuoles in *Coelastrella* cells using Cell Tracker Blue.

**Fig. S7:** Elemental composition of growth media used for *Coelastrella* cells exposed to U.

**Fig. S8:** Growth and photosynthesis of *Coelastrella* cells challenged with U.

**Fig. S9:** PCA of normalized gene expression values for all samples.

**Fig. S10:** Dynamics of differential gene expression during the U stress response in *Coelastrella*.

**Fig. S11:** QPCR analysis of DEGs identified through RNA-seq.

**Fig. S12:** Regulation of genes involved in cell division.

**Fig. S13:** Regulation of genes involved in protein turnover.

**Fig. S14:** Regulation of genes involved in vesicle-mediated transport (A) and autophagy (B).

**Fig. S15:** Regulation of genes and proteins involved in the antioxidant response.

**Fig. S16:** Regulation of genes and proteins involved in photosynthesis.

**Fig. S17:** Regulation of genes involved in chlorophyll synthesis (A) and catabolism (B).

**Movie S1:** 3D reconstructed *Coelastrella* sp. PCV cell grown in mixotrophic conditions.

**Table S1**: **Genomic datasets used in this study**.
HiFi reads are considered for PacBio statistics. *The estimated genome coverage was calculated considering the predictable genome size of *Coelastrella* sp. PCV is 80 Mbp (Karpagam *et al*., 2018; Shetty *et al*., 2021).

| | PacBio Sequel RSII | Illumina NovaSeq 6000 |
|---|---|---|
| Sequencing layout | Single-end long reads | Paired-end (2x150bp) |
| Read bases (bp) | 25,041,797,822 | 14,793,780,456 |
| Number of reads | 1,667,569 | 97,972,056 |
| Average read length (bp) | 15,016 | - |
| Genome coverage* | 313x | 185x |

**Table S2**: **Genome assembly evaluation**.

| | |
|---|---|
| Number of contigs | 407 |
| Total size of contigs (bp) | 119,848,939 |
| Longest contig (bp) | 11,149,823 |
| N50 (bp) | 5,968,012 |
| L50 | 8 |
| GC (%) | 51.71% |

**Table S3**: **Primers used for QPCR analysis**

| Gene ID | Orientation | Sequence 5'>3' |
|---|---|---|
| g5680 (actin) | Forward | TGTTCAACCCCAGCATGGTT |
| | Reverse | TGAGTACTTGCGCTCTGGTG |
| g11267 | Forward | GCAGCAAAGACCCTTGTGTG |
| | Reverse | ACATGATCTGGACTCGACGC |
| g6168 (FTR1) | Forward | AACGCTCGAAGCATCAGTCA |
| | Reverse | ATCCCGGTTATAGCTCCCCA |
| g9648 (FOX1) | Forward | ACCCTGGAACTTGGCTGTTC |
| | Reverse | CCAACTTCACTTTGGGTGCG |
| g13245 | Forward | ACAGCACTCTGAAAGCCCTC |
| | Reverse | GCCAGCTGTGCTAATCCAGA |

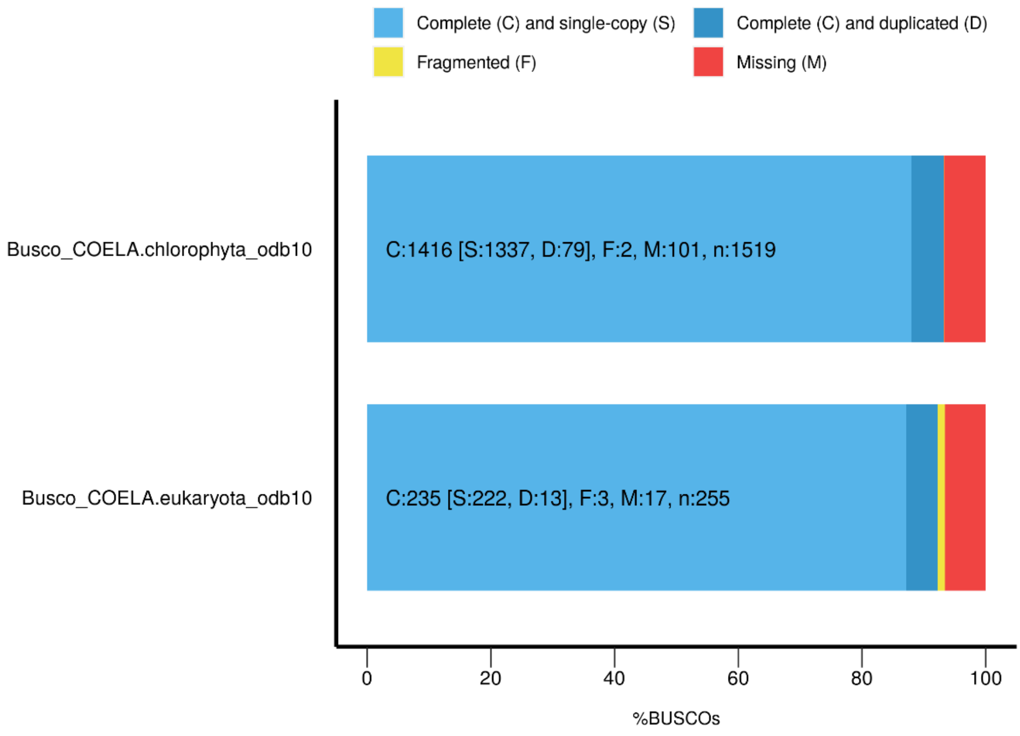


**Figure S1: Assessment of the *Coelastrella* sp. PCV genome completeness using BUSCO.** BUSCO analysis was performed using the Chlorophyta (top) and Eukaryota (bottom) databases. The number of genes in each of the following categories is indicated into brackets: C=complete; S=single copy; D=duplicated; F=fragmented; M=missing; n=total number. The analysis showed 92.2% completeness using Eukaryota markers and 93.2% using Chlorophyta markers.

A

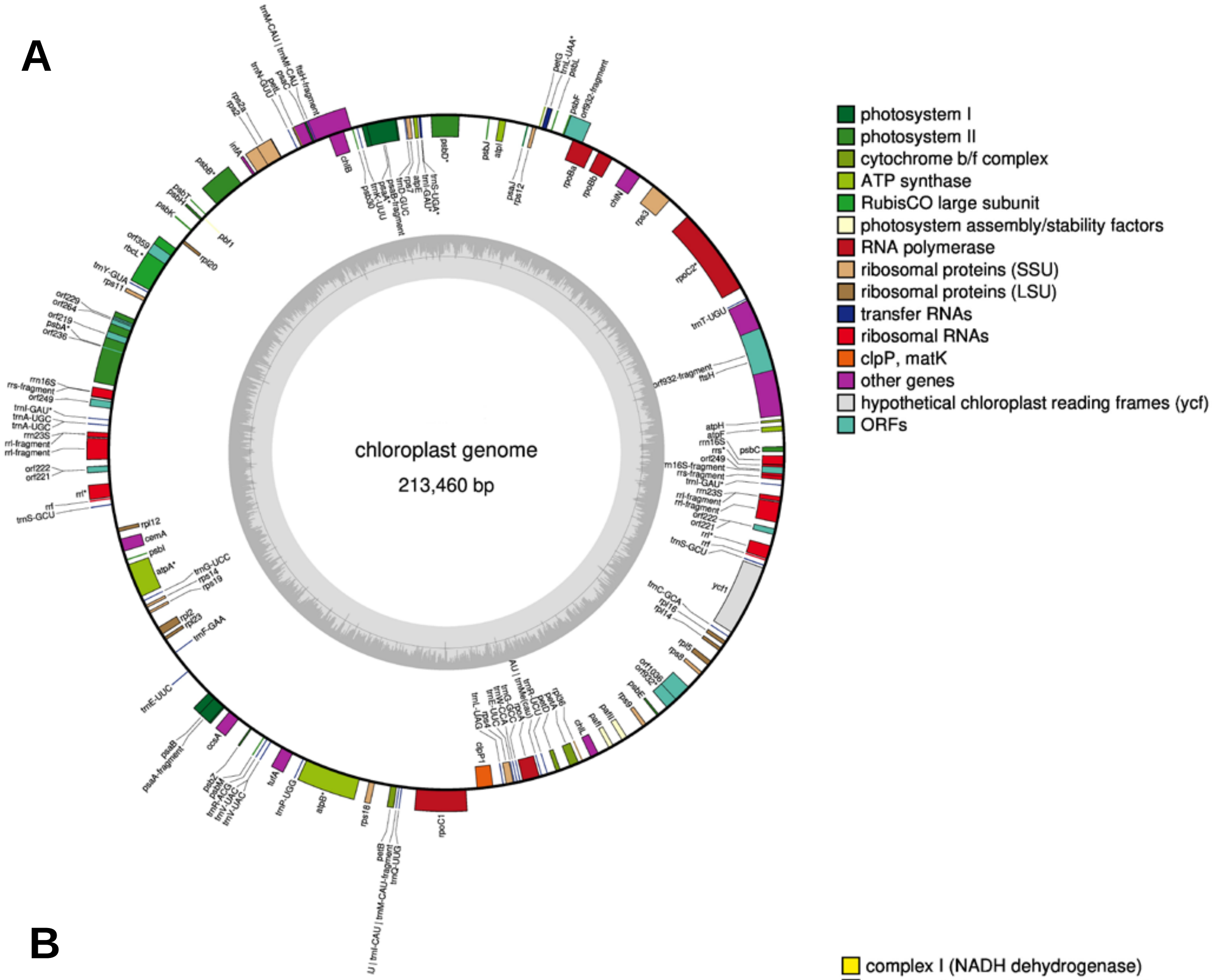


B

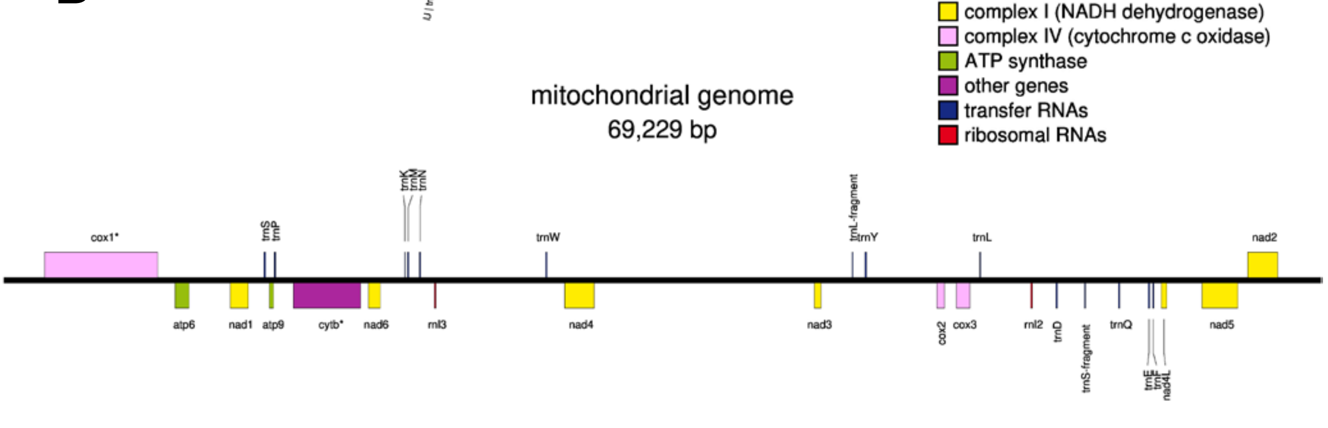


**Figure S2: Structure and annotation of the assembled *Coelastrella* sp. PCV chloroplast (A) and mitochondrial (B) genomes.** The assembled organellar genomes were annotated with the GeSeq online service (Tillich *et al*., 2017). Structure and annotation were produced with OGDRAW (Greiner *et al*., 2019). The chloroplast genome is represented as a typical circular molecule whereas the mitochondrial genome, whose structure is more diverse among green algae, is depicted as linear.

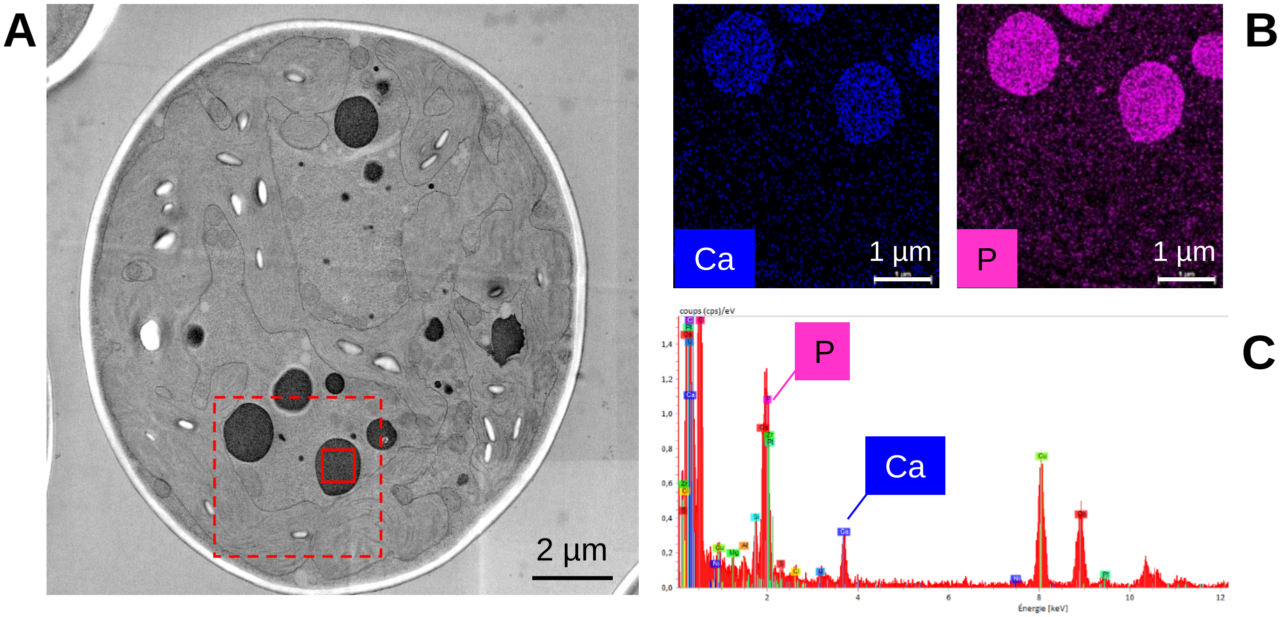


**Figure S3: EDX analysis of acidocalcisomes in *Coelastrella* sp. PCV.**
Cells grown in TAP medium were cryo-fixed, cryo-substituted and analyzed by TEM-EDX. **A** – TEM micrograph of a representative *Coelastrella* cell. **B** – Calcium and phosphorus EDX maps for the area of interest (dotted-line square). **C** – EDX spectrum obtained from an acidocalcisome filled with polyP granules (solid-line square).

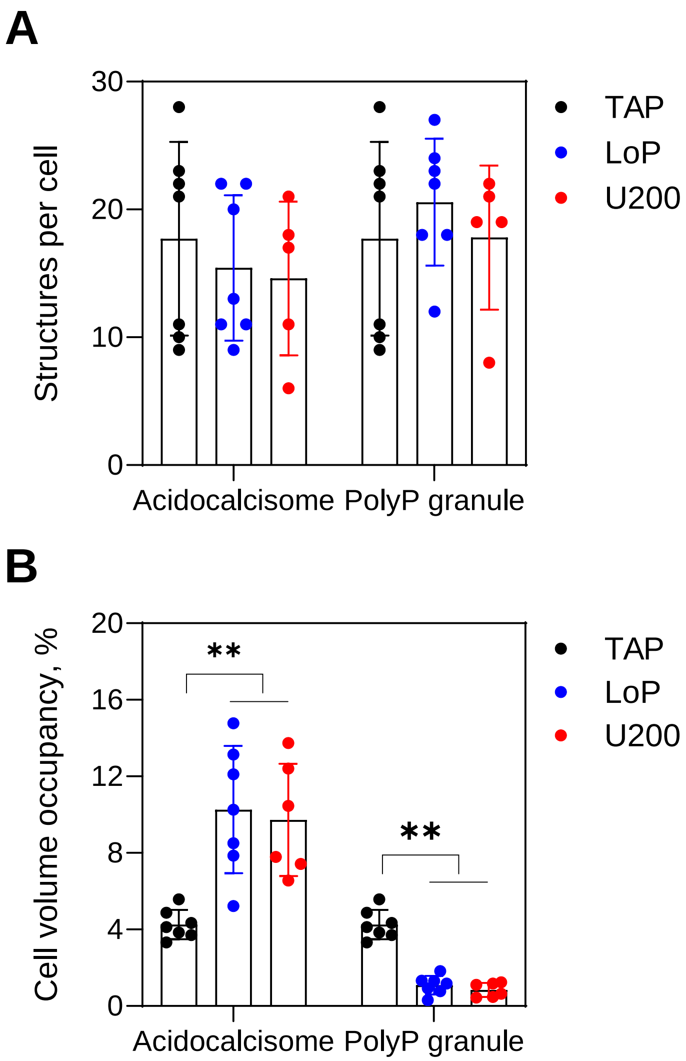


**Figure S4: Morphometric parameters of acidocalcisomes and polyP granules in *Coelastrella* cells challenged with U.** FIB-SEM images were used to measure the number and volume of acidocalcisomes and polyP granules. The maximal diameter of cells and subcellular structures was measured and the corresponding volumes were calculated by approximating these objects as spheres. Measurements have been done for n=6-7 independent cells per condition. Data were analyzed using the Mann-Whitney test. Significance is indicated as p<0.01 (**).

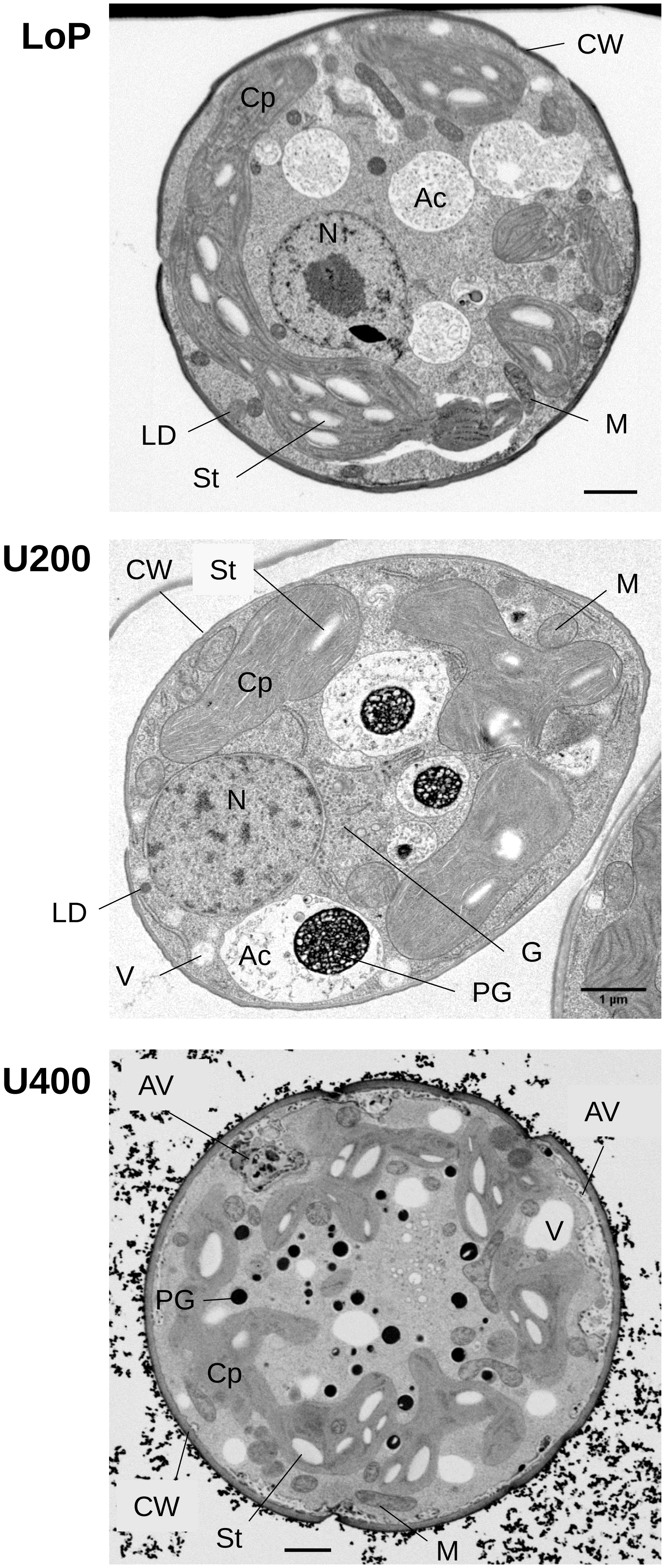


**Figure S5: TEM micrographs of *Coelastrella* in LoP, U200 and U400 conditions.** Cells grown for 24 h in either LoP, U200 or U400 conditions were vitrified and analyzed using FIB-SEM. Selected micrographs are representative of typical cells found in the respective stacks and show the main subcellular structures. Ac, acidocalcisome; Cp, chloroplast; CW, cell wall; G, Golgi; LD, lipid droplet; Mt, mitochondrion; N, nucleus; PG, polyP granule; St, starch granule; autophagic-like vacuoles, AV. Scale bars, 1 µm.

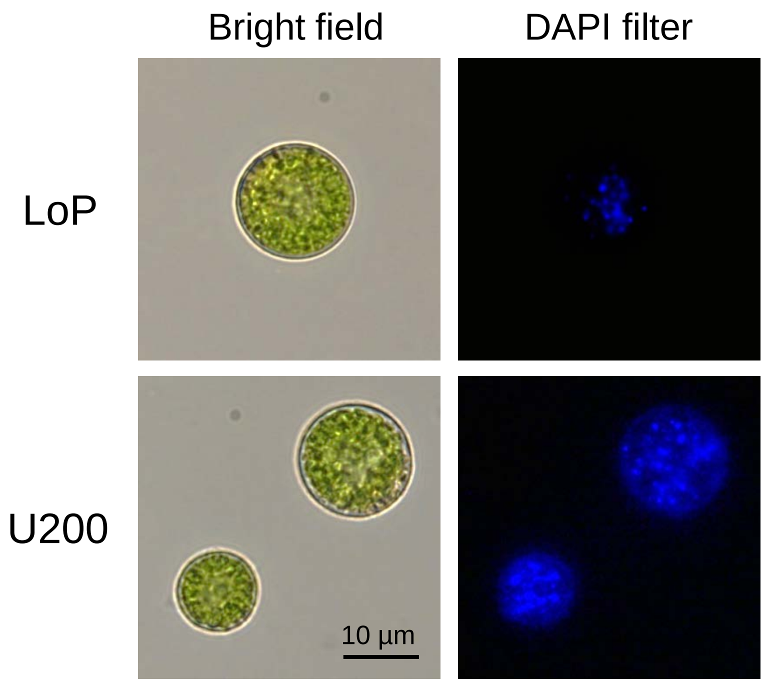


**Figure S6: Observation of vacuoles in *Coelastrella* cells using Cell Tracker Blue.** Cells grown for 24 h in either LoP or U200 were stained using Cell Tracker Blue CMAC, rinced with LoP, and observed by fluorescence microscopy. Cells were imaged using a Zeiss Axioplan 2 microscope operating in bright field or with the DAPI fluorescence filter.

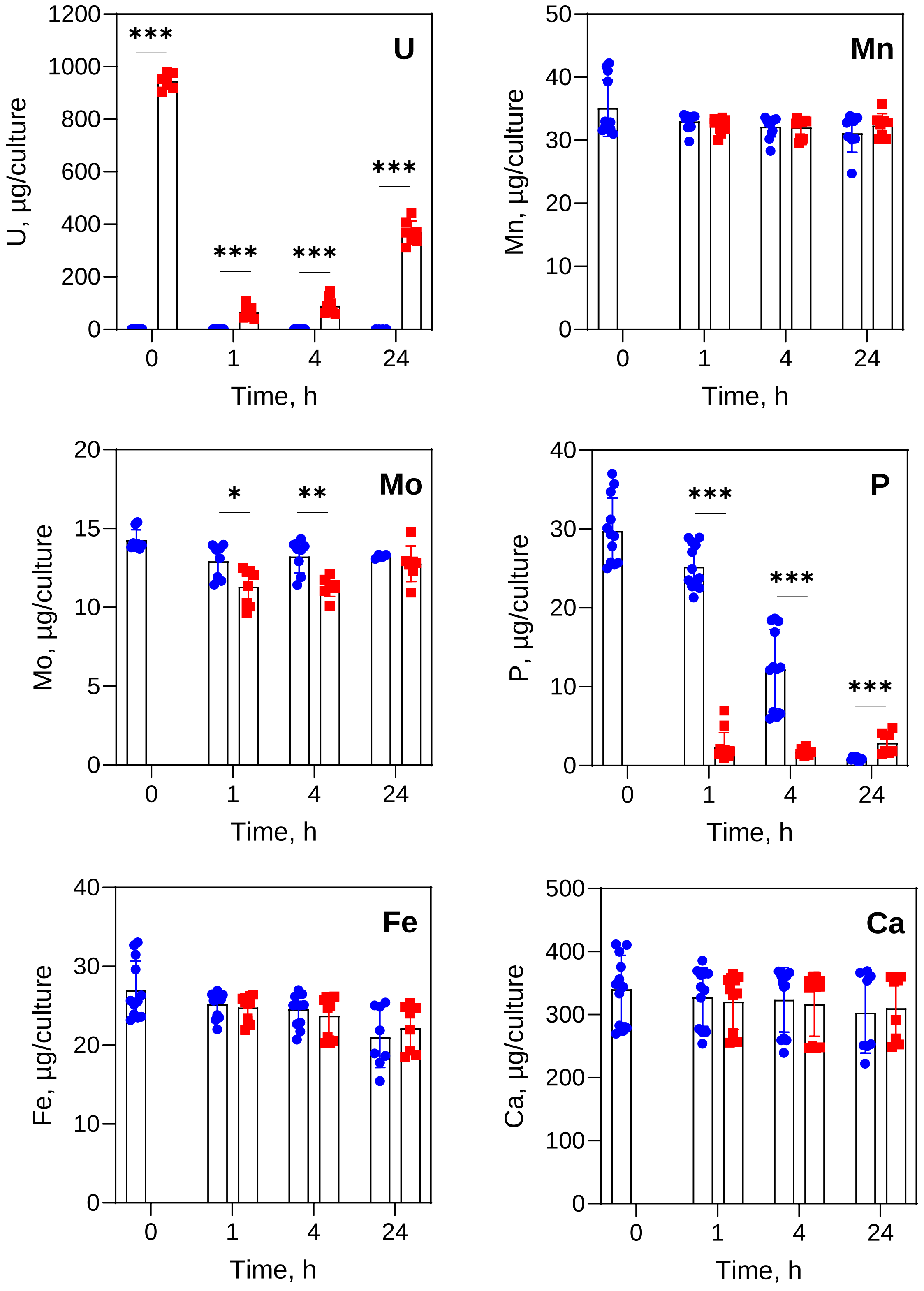


**Figure S7: Elemental composition of growth media used for *Coelastrella* cells exposed to U.** Cells were grown in LoP (●) or U200 (■) for 0, 1, 4, and 24 h and separated from the culture medium by centrifugation. Elemental composition of the culture medium was determined by ICP-MS and is reported as µg of each element per 20-mL culture. Data were analyzed using the Mann-Whitney test. n=7-12 independent cultures per condition. Significance is indicated as $p<0.05$ (*), $p<0.01$ (**), $p<0.001$ (***).

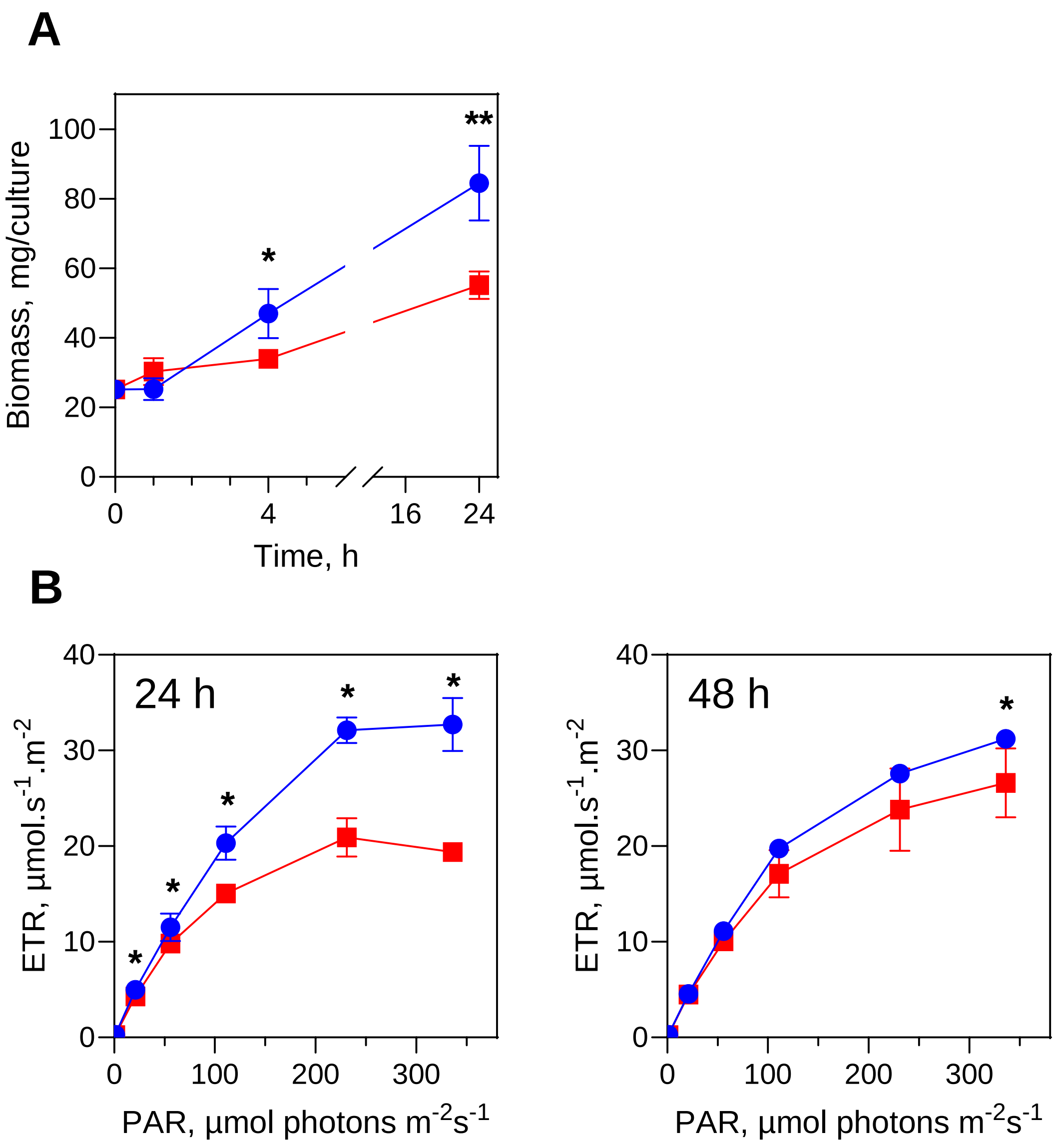


**Figure S8: Growth and photosynthesis of *Coelastrella* cells challenged with U. A -** Cells were grown in LoP (●) or U200 (■) and harvested by centrifugation at different time points. Fresh biomass was determined for each culture (20 mL). **B** - Electron tranfer rate was measured in dark-adapted cells exposed to increasing light intensities after 24 h (left panel) and 48 h (right panel) of growth in LoP (●) or U200 (■). Data are presented as mean ± SD with n=4-5 independent cultures per condition. Statistical analysis was performed using the Mann-Whitney test. Significance is indicated as $p<0.05$ (*), $p<0.01$ (**).

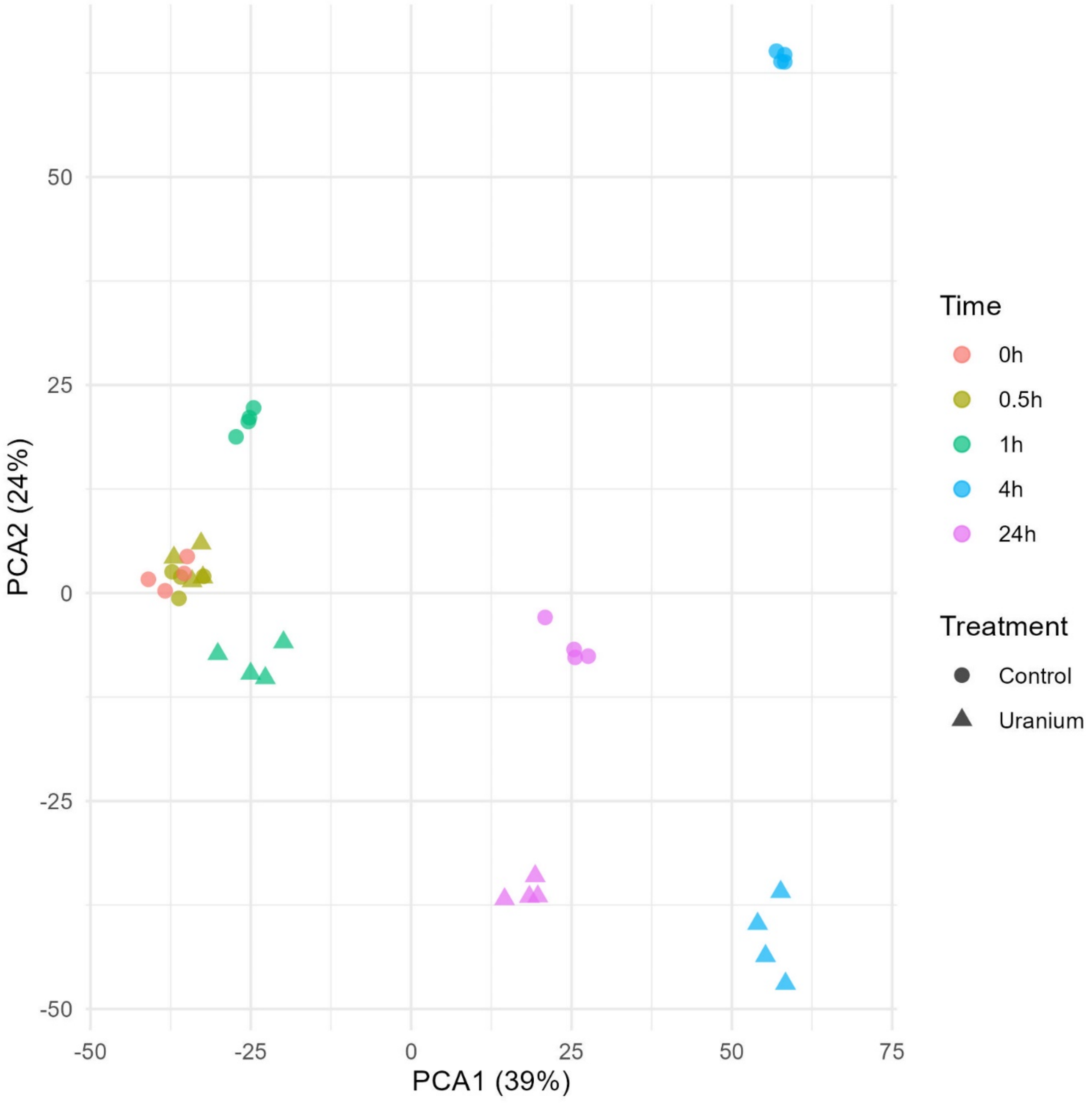


**Figure S9: PCA of normalized gene expression values for all samples.** Raw fragment counts were normalized using the Trimmed Means of M-values method and subsequently filtered and denoised with ARSyNseq. Samples (in quadruplicate) are labeled according to treatment (● control, ▲ uranium) and kinetic time point (color code).

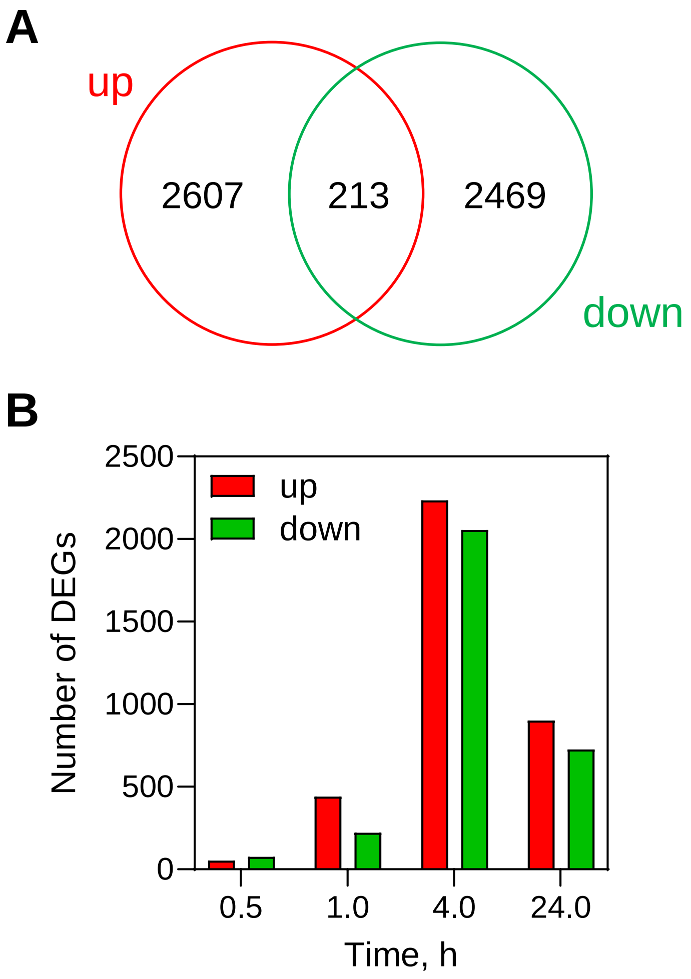


**Figure S10: Dynamics of differential gene expression during the U stress response in *Coelastrella*. A** – Venn diagram showing the overlap of up- and down-regulated genes across the experiment. **B** – Temporal distribution of DEGs throughout the kinetic time course. A total of 5,289 genes were significantly regulated during the stress period, defined by ≥1 FPKM in at least one time point, $|\log_2 FC| \geq 1$ and FDR ≤ 0.01 between control and treated samples at the same time point.

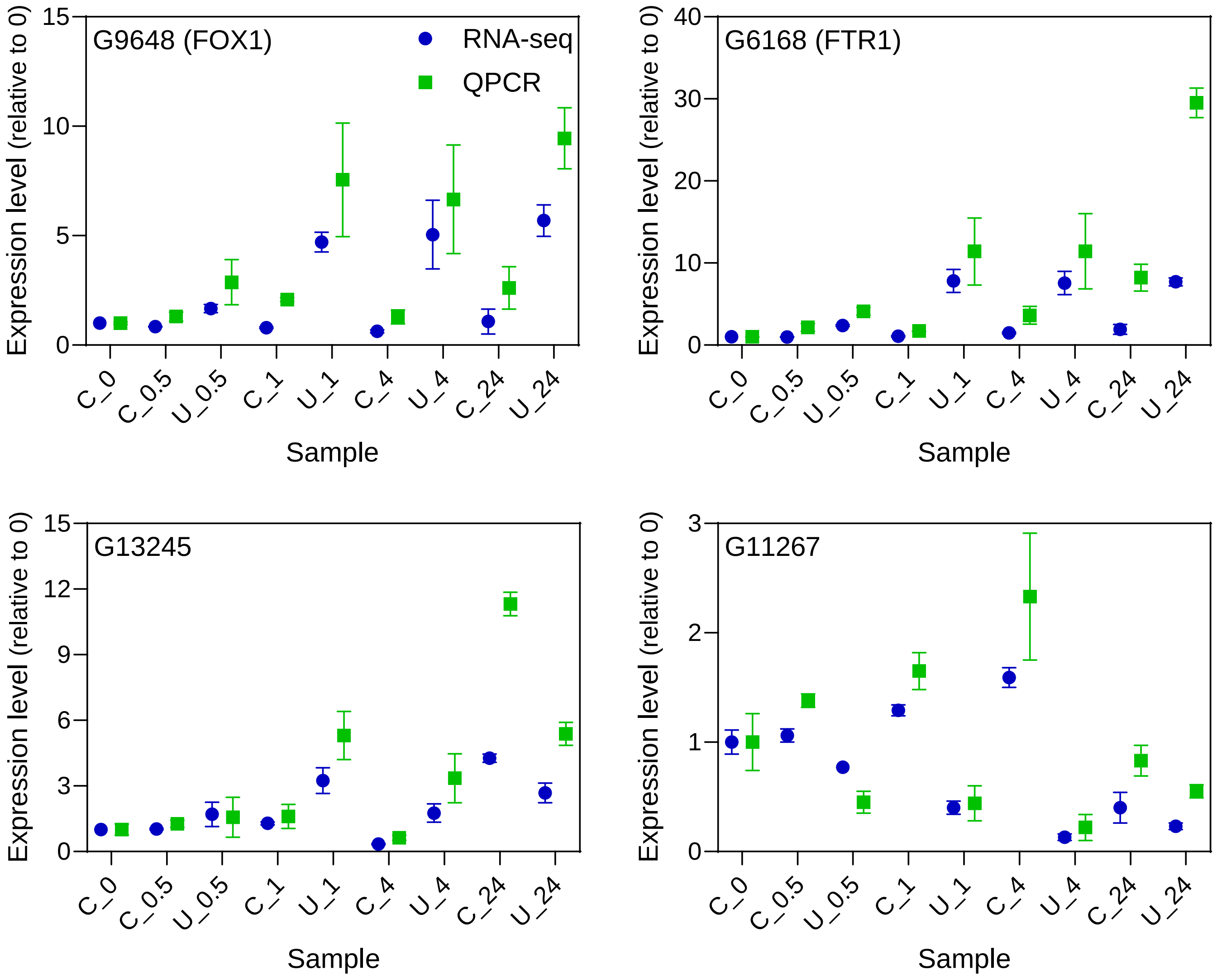


**Figure S11: QPCR analysis of DEGs identified through RNA-seq.** The expression of four DEGs was analyzed by QPCR using total RNAs from the RNA-seq experiment. RNA-seq expression levels (●) are mean ± SD of four replicates, while QPCR data (■) are from two of these replicates. Sample are labeled according to treatment (C, control; U, uranium) and time point. Primers used for QPCR analysis are shown in Table S3.

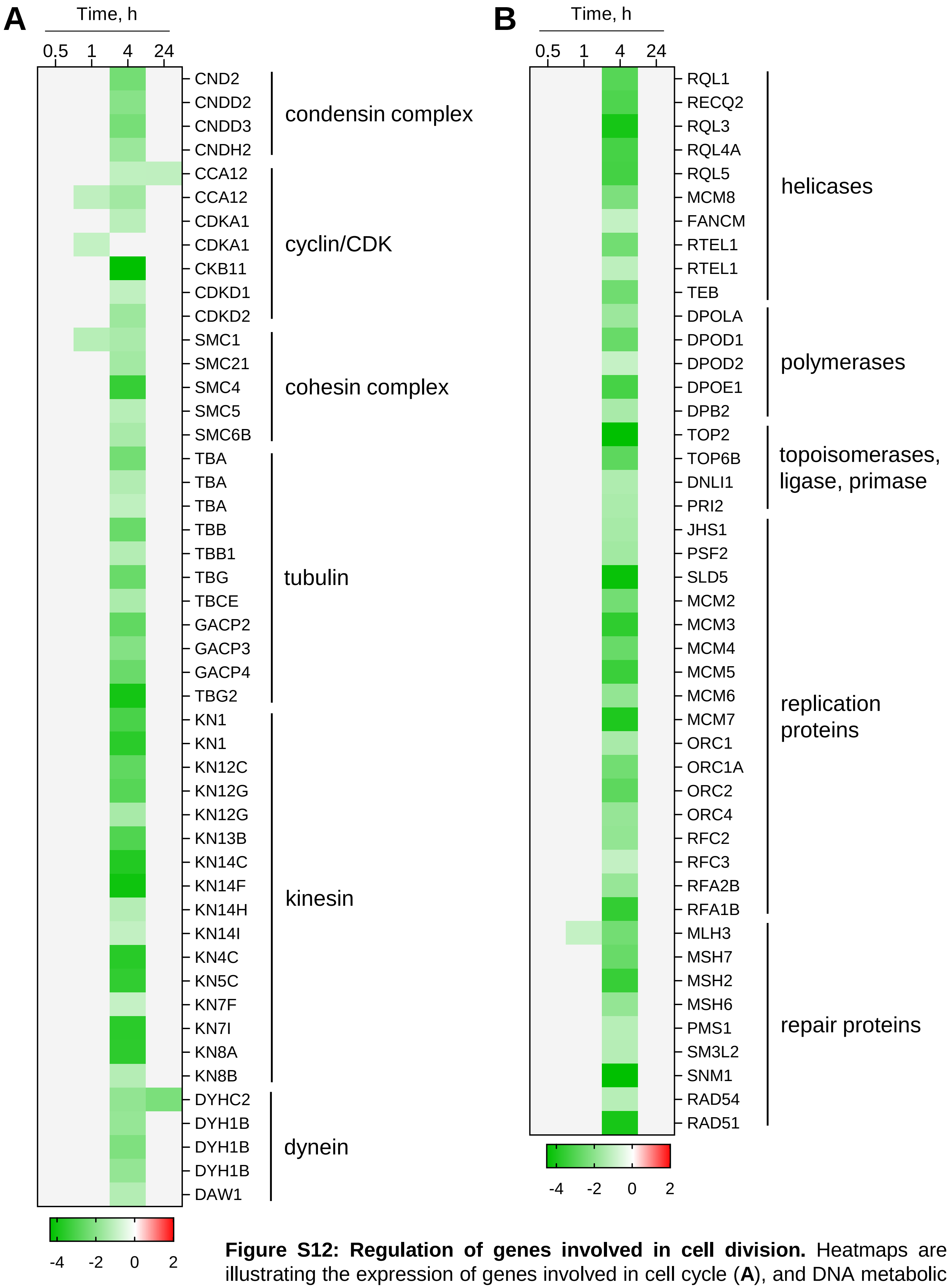


**Figure S12: Regulation of genes involved in cell division.** Heatmaps are illustrating the expression of genes involved in cell cycle (**A**), and DNA metabolic process (**B**). Genes showing significant changes in expression in response to U stress (FPKM ≥1, |$\log_2$FC| ≥1, FDR ≤0.01) are colored red (upregulated) or green (downregulated). Non-significant changes are shown in light grey. Expression values are displayed on a $\log_2$ scale.

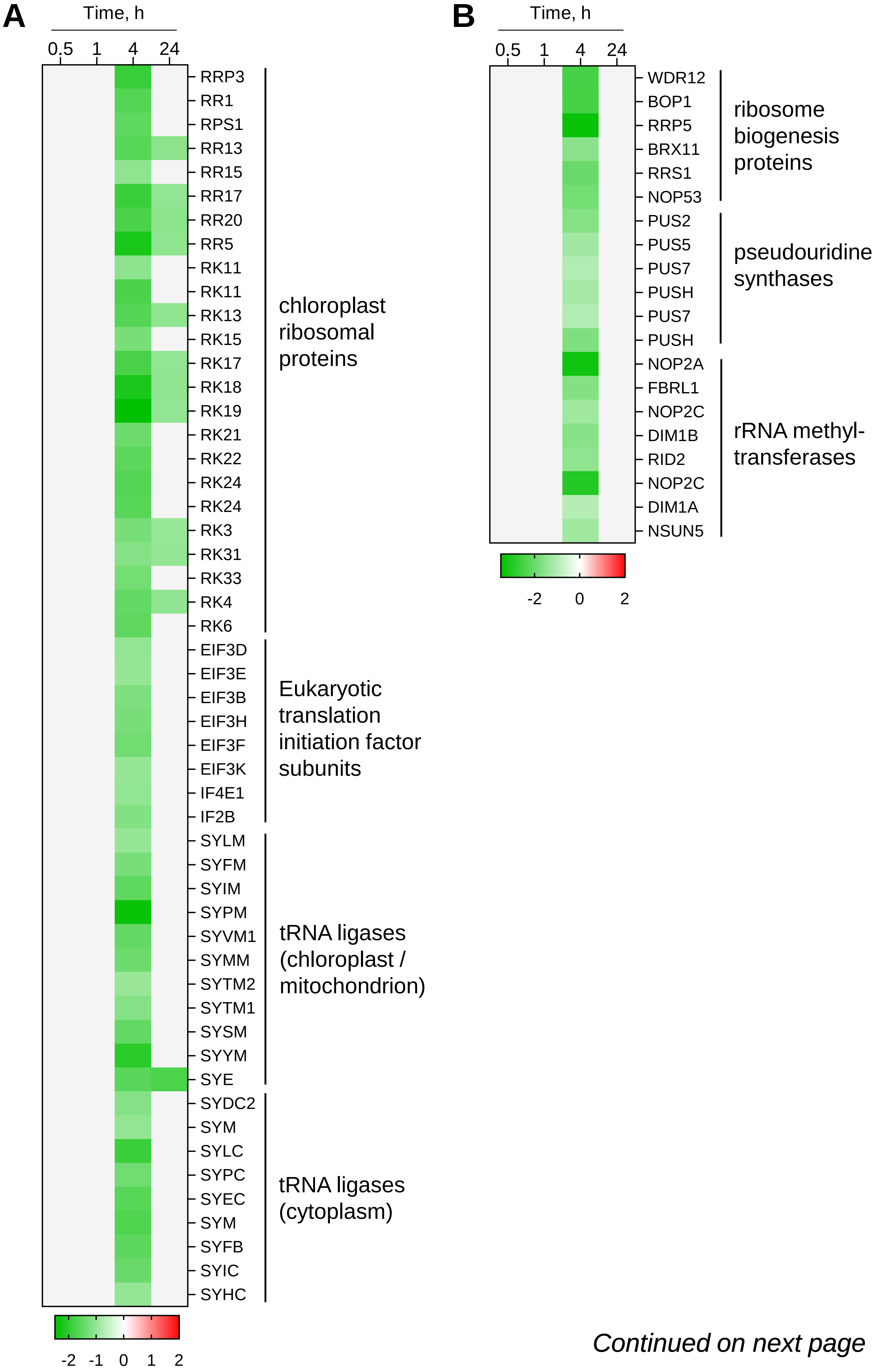


*Continued on next page*

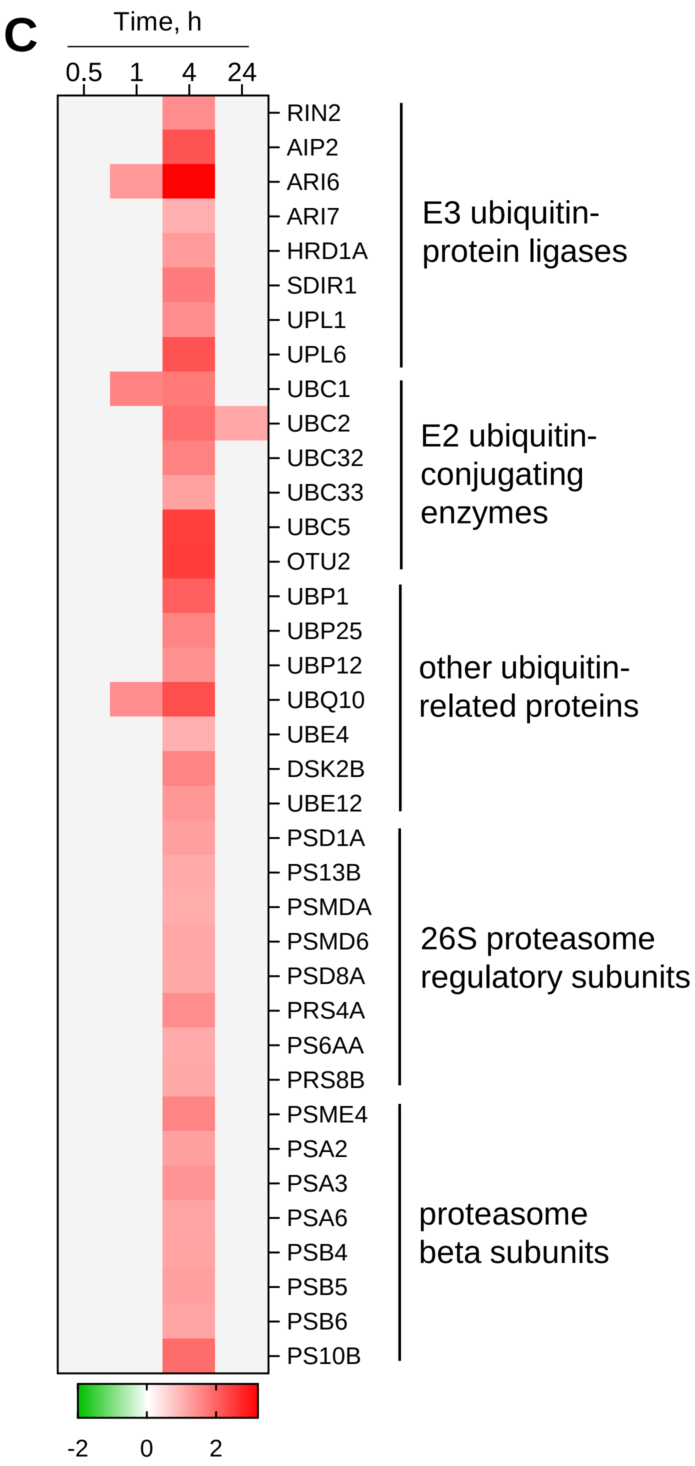


**Figure S13: Regulation of genes involved in protein turnover.** Heatmaps are illustrating the expression of genes involved in translation (**A**), ribosome biogenesis (**B**), and ubiquitin-dependent protein catabolism (**C**). Genes showing significant changes in expression in response to U stress (FPKM ≥1, $|\log_2 FC|$ ≥1, FDR ≤0.01) are colored red (upregulated) or green (downregulated). Non-significant changes are shown in light grey. Expression values are displayed on a $\log_2$ scale.

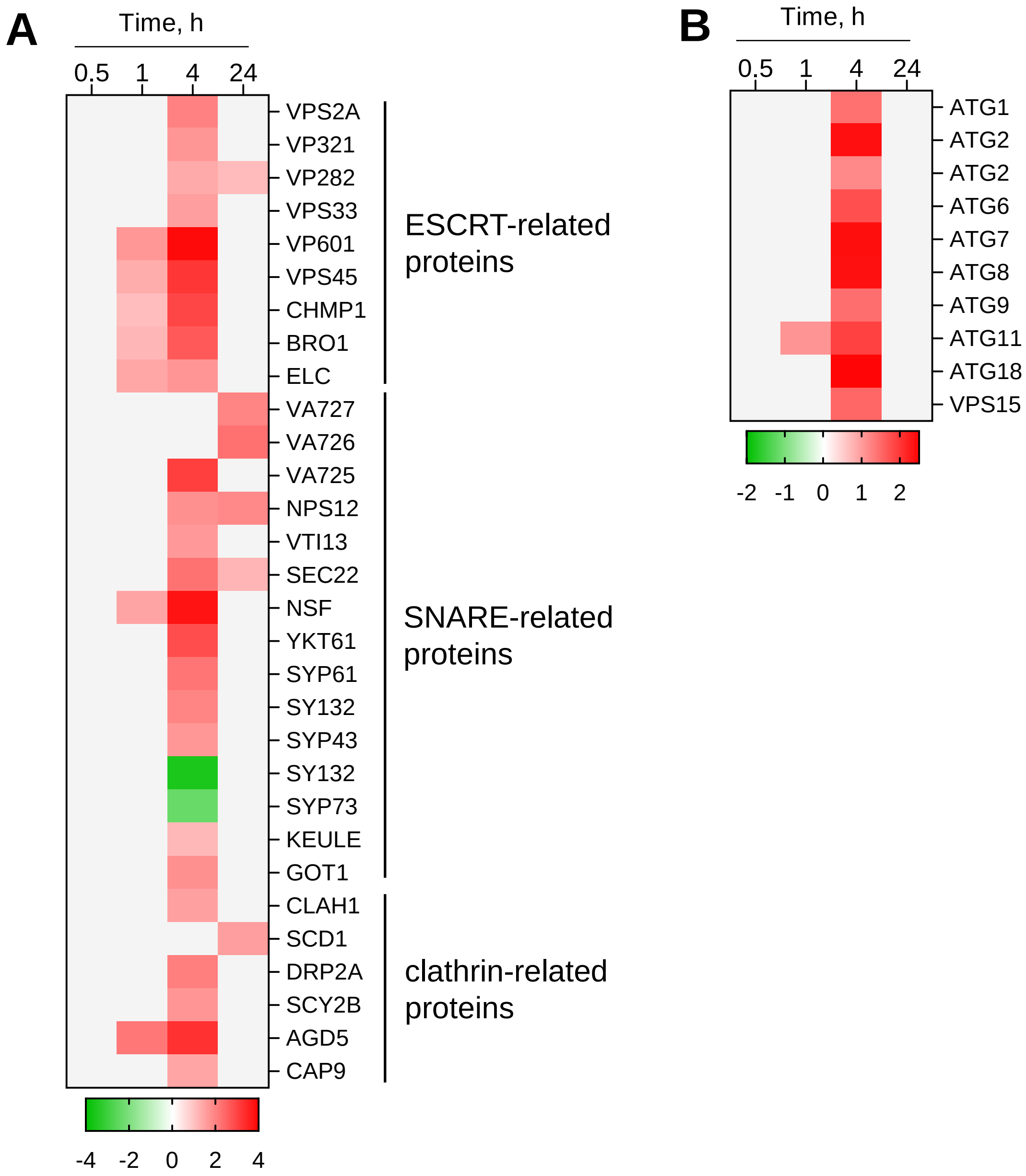


**Figure S14: Regulation of genes involved in vesicle-mediated transport (A) and autophagy (B).** Genes showing significant changes in expression in response to U stress (FPKM ≥1, $|\log_2 FC|$ ≥1, FDR ≤0.01) are colored red (upregulated) or green (downregulated). Non-significant changes are shown in light grey. Expression values are displayed on a $\log_2$ scale.

A

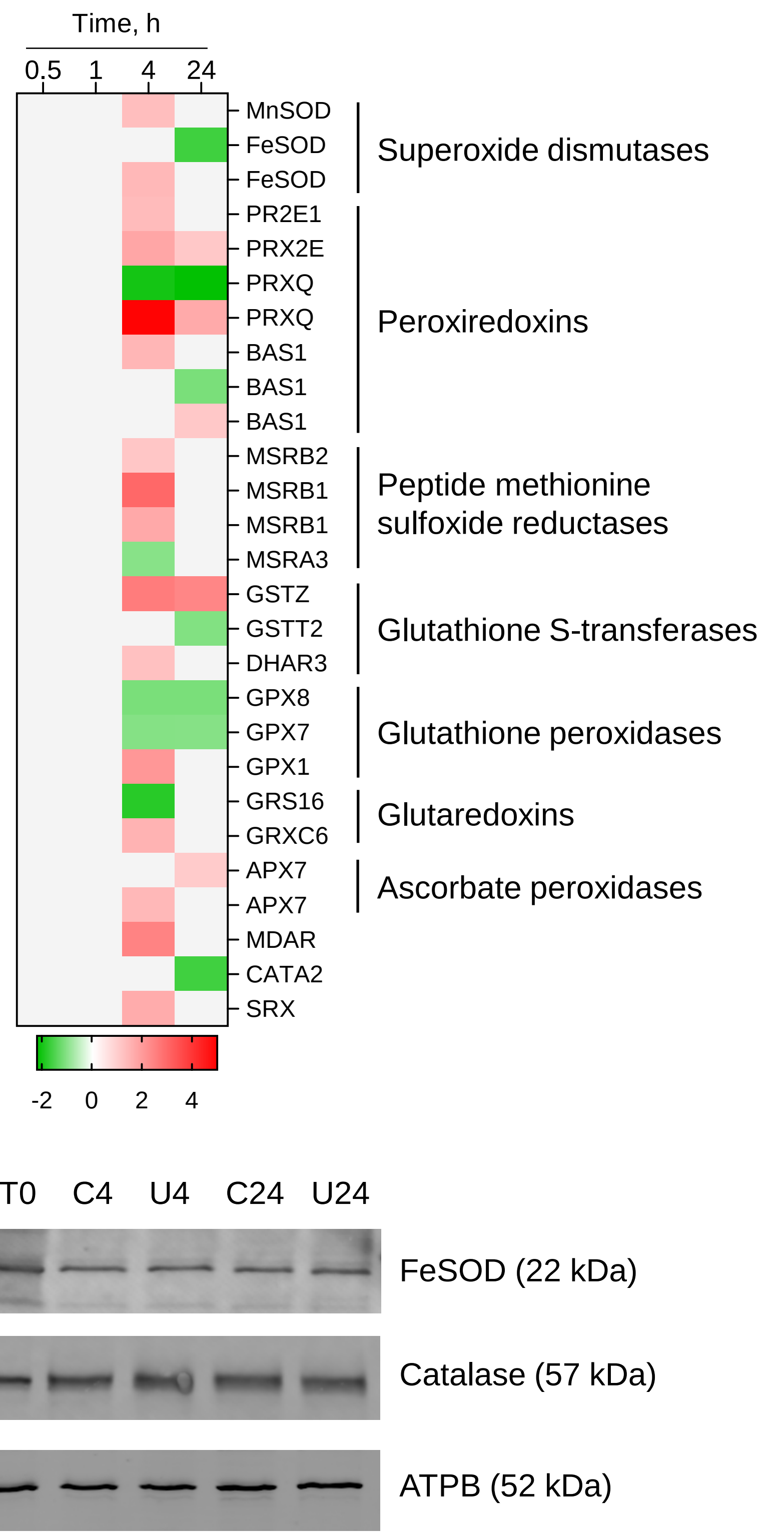


B

**Figure S15: Regulation of genes and proteins involved in the antioxidant response. A –** Heatmap illustrating the expression of genes involved in the antioxidant response. Genes showing significant changes in expression in response to U stress (FPKM ≥1, $|log_2FC|$ ≥1, FDR ≤0.01) are colored red (upregulated) or green (downregulated). Non-significant changes are shown in light grey. Expression values are displayed on a $log_2$ scale. **B** - Immunoblot analysis of Fe-dependent SOD and catalase. Total protein extracts were prepared from *Coelastrella* cells at time zero (T0), 4 h and 24 h in control (C) or U200 (U) conditions. FeSOD, catalase, and ATPB (loading control) were probed using specific antibodies.

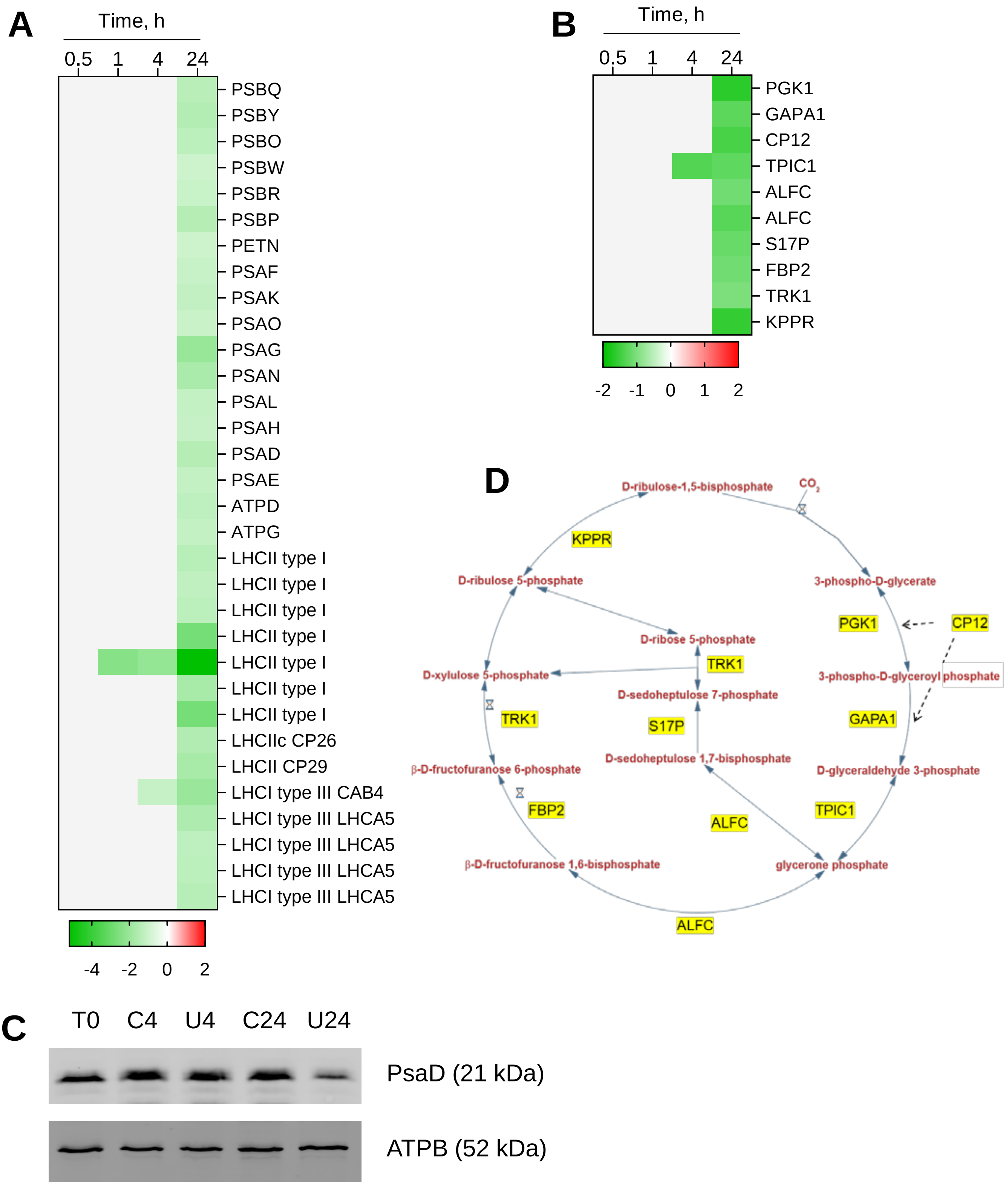


**Figure S16: Regulation of genes and proteins involved in photosynthesis.** Heatmaps are illustrating the expression of genes involved in photosynthesis light reactions (**A**) and the Calvin cycle (**B**). Genes showing significant changes in expression in response to U stress (FPKM ≥1, $|log_2FC|$ ≥1, FDR ≤0.01) are colored red (upregulated) or green (downregulated). Non-significant changes are shown in light grey. Expression values are displayed on a $log_2$ scale. **C** - Immunoblot analysis of PsaD and ATPB. Total protein extracts were prepared from *Coelastrella* cells at time zero (T0), 4 h and 24 h in control (C) or U200 (U) conditions. PsaD and ATPB were probed using specific antibodies. **D** – Representation of the Calvin cycle highlighting steps regulated at the gene level. Pathway overview was obtained from the Plant Metabolic Network (https://plantcyc.org/).

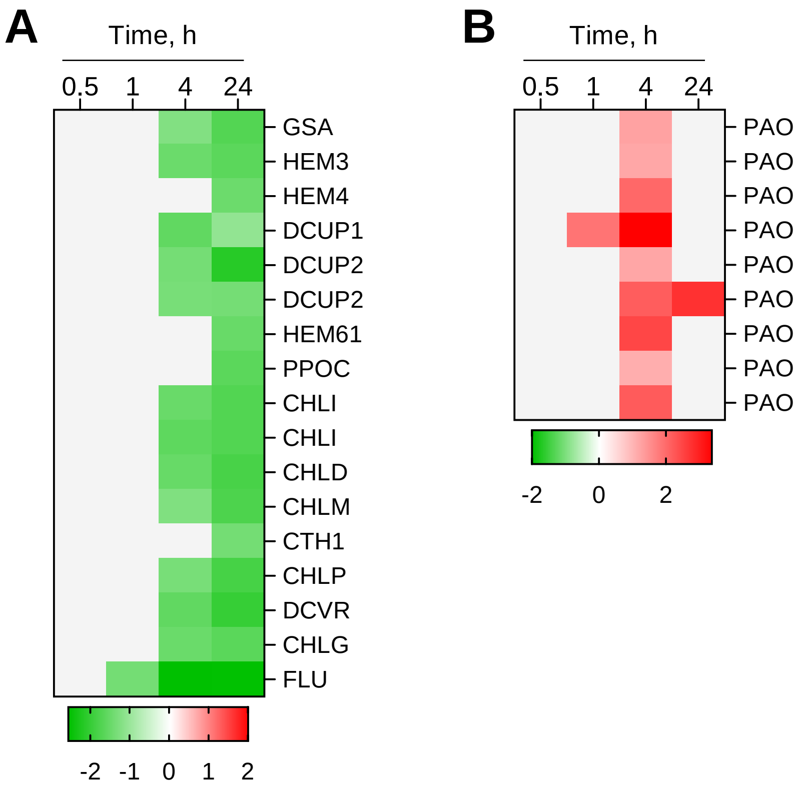


**Figure S17: Regulation of genes involved in chlorophyll synthesis (A) and catabolism (B).** Genes showing significant changes in expression in response to U stress (FPKM ≥1, $|\log_2 FC|$ ≥1, FDR ≤0.01) are colored red (upregulated) or green (downregulated). Non-significant changes are shown in light grey. Expression values are displayed on a $\log_2$ scale.
